# Nitroxoline evidence Amoebicidal Activity against *Acanthamoeba castellanii* through DNA damage and the stress response pathways

**DOI:** 10.1101/2024.09.29.615638

**Authors:** Lijun Chen, Wei Han, Meng Feng, Wenwen Jing, Qingtong Zhou, Xunjia Cheng

## Abstract

*Acanthamoeba castellanii* is a widespread unicellular eukaryote found in diverse environments, including tap water, soil, and swimming pools. It is responsible for severe infections, such as *Acanthamoeba* keratitis and granulomatous amebic encephalitis, particularly in individuals with immunocompromisation. The ability of protozoans to form dormant cysts complicates treatment, as current therapies are ineffective against cyst stages and suffer from poor specificity and side effects. Nitroxoline, a quinoline derivative with well-established antibacterial, antifungal, and antiviral properties, is a promising therapeutic candidate. This study aimed to elucidate cellular signalling events that counteract the effects of nitroxoline. In this study, nitroxoline significantly reduced the viability of *A. castellanii* trophozoites in a dose- and time-dependent manner, inducing morphological changes and apoptosis. Transcriptomic analysis revealed substantial alterations in gene expression, including enrichment of metabolic pathways, DNA damage responses, and iron ion binding. Nitroxoline treatment upregulated genes involved in DNA repair and oxidative stress response while regulating genes in the methionine and cysteine cycles. It also decreased the mitochondrial membrane potential, H₂S production, and total iron amount in *A. castellanii*. Bioinformatics analyses and molecular docking studies suggest direct interactions between nitroxoline and *A. castellanii* proteins. Our research provides a comprehensive molecular map of the response of *A. castellanii* to nitroxoline, revealing significant changes in gene expression related to the stress response and metabolic pathways. These findings underscore the potential of nitroxoline as a potent anti-*Acanthamoeba* agent, offering new insights into its mechanism of action and paving the way for effective therapeutic strategies.

## 1. Introduction

*Acanthamoeba castellanii* is a cosmopolitan unicellular eukaryotic organism commonly found in tap water, swimming pools, soil [1], cooling towers, and drinking water networks. Protozoa are opportunistic amoebae with a widespread distribution worldwide [2,3]. In humans, *Acanthamoeba* species are the most common causal organisms [4], cause sight-threatening keratitis named *Acanthamoeba* keratitis (AK) [5], and in patients with immunocompromisation, it also results in severe invasive infections, such as granulomatous amoebic encephalitis and chronic infectious ulcers [6–8]. *Acanthamoeba* can cause AK in immunocompetent individuals, and its distribution is limited to the eye, where it can cause blind keratitis. The annual incidence of AK is approximately 1.2 million in Western countries [9,10].

*Acanthamoeba* also has the innate ability to survive under harsh environmental conditions and lethal drug exposure without genetic resistance by transiently stopping their growth, slowing their metabolism, and eventually shifting to dormant and inactive cysts. A conceptually compelling but sparsely investigated approach for identifying useful potentiators involves disrupting the general defence systems that protect pathogens from diverse antibiotics. Currently, no effective anti-*Acanthamoeba* drugs are available. Therefore, innovative anti-*Acanthamoeba* drugs are urgently required. Most current anti-*Acanthamoeba* drugs target the trophozoite-stage parasites. Although cyst-stage parasites do not cause clinical symptoms, drugs that inhibit these two stages are essential for preventing epidemics and protecting vulnerable populations, owing to the increase in anti-*Acanthamoeba* drug resistance. Chemotherapy for *Acanthamoeba* infections remains challenging due to incomplete knowledge of its complicated pathophysiology. In cases of infection, the treatment regimen is often ineffective owing to delayed diagnosis, poor specificity, and side effects. However, this route is lengthy, with many hurdles and failures, including target specificity, poor pharmacodynamics, pharmacokinetics, and toxicity.

Considering the limited number of anti-*Acanthamoeba* agents, screening of potential anti-*Acanthamoeba* agents from natural products or approved drugs is warranted. Nitroxoline, a quinoline derivative (8-hydroxy-5-nitroquinoline), is a historically used oral antibacterial drug [11] that is structurally distinct from other antibiotics. In addition to its antitumor effects, it is active against numerous human pathogens, including multidrug-resistant bacteria, fungi, and viruses [12–15]. Its antibacterial properties are based on the chelation of divalent cations, influencing cell membranes, and inhibiting intracellular enzymes such as RNA polymerase [16]. From a mechanistic standpoint, nitroxoline partially interferes with the phosphatidylinositol 3-kinase/AKT/mammalian target of rapamycin and rapidly accelerated fibrosarcoma/mitogen-activated protein kinase/extracellular signal-regulated kinase host cell signalling pathways [17].

Recent studies have also indicated that nitroxoline is a promising drug against free-living pathogenic amoebas, including *Naegleria fowleri*, *Balamuthia mandrillaris*, and *Acanthamoeba* species [18–20]. Although transcriptomic and proteomic analyses have been performed on these free-living amoebae, the signalling pathways involved in their stress responses remain limited. Furthermore, our knowledge of the transcriptomic landscape during nitroxoline treatment remains limited. This study aimed to elucidate the cellular signalling events that counteract the effects of nitroxoline.

## 2. Materials and methods

### 2.1. Amoeba cultivation

*Acanthamoeba castellanii* (American Type Culture Collection (ATCC) 30011) was obtained from the ATCC. Following established protocols, trophozoites were axenically cultured in a peptone–yeast–glucose medium. This medium comprised 10 g of protease peptone, 10 g of yeast extract, 10 g of glucose, 5 g of NaCl, 0.95 g of L-cysteine, 3.58 g of Na_2_HPO_4_·12H_2_O, and 0.68 g of KH_2_PO_4_ per litre of deionised distilled water. The culture was incubated at 26°C, and trophozoites, harvested during the late log phase after 48 h of subculture, were used for subsequent analyses.

### 2.2. Cell viability assay and IC50 determination

Obtained from TargetMol (Wellesley Hills, MA, USA), nitroxoline was dissolved in 100% dimethyl sulfoxide (DMSO; Sigma-Aldrich, St. Louis, MO, USA) and prepared at 100 mM concentrations. *A. castellanii* trophozoites were seeded at the log growth phase (1 × 10^4^ cells per well) in a 96-well white microplate (Eppendorf, Hamburg, Germany). Incubation with medium containing varied nitroxoline concentrations ranging from 0 to 100 μM was performed, and the plates were incubated at 26°C for 24 h, 48 h, and 72 h and observed under an inverted microscope (Olympus). Cell viability assay results, derived from ≥3 wells per condition and compared with diluted DMSO, were evaluated using CellTiter-Glo (Promega, Madison, WI, USA) following the manufacturer’s guidelines. Subsequently, 100 μL of CellTiter-Glo reagent was introduced per well, mixed for 2 min on an orbital shaker, and incubated at room temperature for 10 min. Luminescence was recorded on a modular multimode microplate reader (BioTek Synergy H1), and the IC50 values of the compound and growth curves were generated using GraphPad Prism 8. Each experiment was conducted thrice.

### 2.3. Flow cytometric analysis

Trophozoite apoptosis and reactive oxygen species (ROS) were assessed using a PE Annexin V apoptosis detection kit I (BD Biosciences, San Jose, CA, USA) and a ROS assay kit (Dojindo Molecular Technologies, Inc., Kumamoto, Japan), respectively, following the manufacturer’s instructions. *A. castellanii* trophozoites (2 × 10^6^ cells/flask) were cultured with various nitroxoline concentrations for 24 h in 12.5 cm^2^ cell culture flasks (Biofil). Trophozoites were then detached from the flasks, collected in transparent centrifuge tubes (15 mL), and centrifuged at 800 × g for 5 min. These trophozoites (1 × 10^6^ cells/tube) underwent two cold Hanks’ balanced salt solution (HBSS) washes and were respectively resuspended in 100 μL of 1× binding buffer containing 5 μL of PE-Annexin V and 500 μL of ROS working solution containing 0.5 μL of highly sensitive DCFH-DA dye. Subsequently, the trophozoites for apoptosis detection were incubated for 15 min in darkness at room temperature, and we added 400 μL of 1× binding buffer after incubating. The trophozoites for ROS detection were incubated for 30 min in darkness at room temperature, washed twice with HBSS, and resuspended with 500 μL of HBSS. Fluorescence-activated trophozoite sorting was performed using a FACSAria instrument (BD Biosciences, San Jose, CA, USA) with a 488-nanometer argon excitation laser. Analysis gates were defined using untreated amoebae, and FlowJo 10.8.1 software (FlowJo LLC, Ashland, OR, USA) was used for data analysis.

### 2.4. RNA-sequencing (RNA-seq)

Total RNA was isolated from each thymic sample using an RNA mini kit (Qiagen, Hilden, Germany). RNA quality was examined using gel electrophoresis and Qubit (Thermo, Waltham, MA, USA). For RNA sequencing, the RNA samples from three biological replicates at 24 h were separated into three independent pools, each containing two or three distinct samples in equal amounts. The raw data were subsequently analyzed as previously described [21].

### 2.5. Real-time quantitative polymerase chain reaction (RT-qPCR)

Total RNA was isolated using the RNeasy Plus Mini Kit (74134; QIAGEN, Hilden, Germany), and cDNA was synthesised using the PrimeScript™ II 1st Strand cDNA Synthesis Kit (6210A; Takara Bio, China). The reactions were performed in a 96-well plate using SYBR Premix Ex Taq (Takara, Dalian, Liaoning, China) with primers targeting *A. castellanii GNMT*, *CBS*, *GSR*, *PARP*, *MRE11*, *RAD50*, *RAD51*, *DNA2*, *FEN1, APE1,* and *DMT1*. Primers for these genes, sourced from published sequences, were used for RT-qPCR on an ABI 7500 RT-PCR system (Applied Biosystems, Foster City, CA, USA). The 2-^△△Ct^ method was used to calculate the relative expression of each primer between the control and treatment groups, with values normalised to the 18s reference housekeeping gene.

### 2.6. DNA damage determination

Genomic DNA was extracted using the DNeasy Blood & Tissue Kit (Catalogue No. 69506; QIAGEN), following the manufacturer’s protocol. DNA concentrations were quantified using a BioPhotometer® D30 (Eppendorf). Trophozoite DNA damage was assessed using the DNA Damage Quantification Kit AP Site Counting (Dojindo, catalogue DK02). DNA was dissolved in TE buffer (10 mM Tris [pH 7.5] and 1 mM EDTA) at 100 μg/mL, and 10 μL of purified genomic DNA was mixed with 10 μL of aldehyde reactive probe (ARP) solution in a MaxyClear microcentrifuge tube (1.5 mL; Corning, NY, USA) and incubated for 1 h at 37°C. Each ARP-labelled DNA was purified with a filtration tube, and DNA concentration was measured using the BioPhotometer® D30. Purified ARP-derived DNA samples were diluted to 2.25 μg/mL in TE buffer, and 90 μL of the ARP-labelled DNA solution was diluted with 310 μL of TE buffer. Thereafter, 60 μL of the diluted solution and 100 μL of DNA binding solution were added to the provided DNA high-binding plate well, incubated overnight at room temperature, and microwells were washed five times in 250 μL of 1× wash buffer. Subsequently, 150 μL of diluted horseradish peroxidase-streptavidin-enzyme conjugate (1:4000 dilution in 1× wash buffer) was added to each well, incubated for 1 h at 37°C, and microwells were washed five times in 250 μL of 1× wash buffer. After adding 100 μL of substrate solution, microwells were incubated at 37°C for 1 h. Absorbance at an optical density (OD) of 650 nm was immediately measured using a modular multimode microplate reader (BioTek Synergy H1).

### 2.7. Mitochondrial membrane potential (MMP) assay

MMP was evaluated using the MT-1 MitoMP detection kit (Dojin Kagaku) following the manufacturer’s protocol. Treated trophozoites (1×10^6^) were incubated with MT-1 working solution (v:v, 1:1000) for 30 min at 26℃. After treatment with the MT-1 working solution, the trophozoites were washed twice with HBSS and incubated in an imaging buffer. Live trophozoite suspensions were placed on glass slides using a Cytospin 4 cytocentrifuge (Thermo Fisher Scientific) and mounted on coverslips. The fluorescence images were obtained using a Leica TCS SP8 microscope, and the fluorescence (λEx = 488 nm and λEm = 580 nm) was measured using a modular multimode microplate reader (BioTek Synergy H1). Images were quantified using ImageJ software (National Institutes of Health, Bethesda, MD, USA).

### 2.8. Measurement of H_2_S production in trophozoites

Following a 24-h treatment, trophozoites (1×10^6^) were washed twice with 1× phosphate-buffered saline (PBS) and incubated for 1 h at 26°C in 500 μL of HBSS containing 100 μM of 7-azido-4-methylcoumarin (AzMC) fluorescent probe (Sigma), which selectively reacts with H_2_S to form a fluorescent compound. Fluorescent AzMC signal acquisition (λEx = 365 nm and λEm = 450 nm) was performed on a modular multimode microplate reader (BioTek Synergy H1), and the fluorescence images were obtained using a Leica TCS SP8 microscope.

### 2.9. Intracellular iron assay

The amount of iron (Fe^2+^ and Fe^3+^) was monitored using an iron assay kit (Dojindo Laboratories Co., Ltd., Kumamoto, Japan). Following a 24-h treatment, trophozoites (2×10^7^) were washed twice with 1× PBS, and 1 mL of assay buffer was added to the kit. The suspension was ultrasonicated for 20 min and centrifuged at 16,000 ×g for 15 min. After following the manufacturer’s instructions, the standard and samples (n = 3 for each group) were incubated at 37°C for 1 h. Absorbance at OD 593 nm was measured using a modular multimode microplate reader (BioTek Synergy H1).

### 2.10. Identifying nitroxoline target homologues in *A. castellanii* proteome

In the Research Collaboratory for Structural Bioinformatics (RCSB) Protein Data Bank (PDB), two entries with nitroxoline as the structural ligand were identified: 3AI8 (Gene: CTSB, UniProt ID: P07858) and 5Y1Y (Gene: BRD4, UniProt ID: O60885) [22,23]. To identify proteins homologous to these two known nitroxoline targets in *A. castellanii*, the complete proteome dataset for *A. castellanii* (Proteome ID: UP000011083) was retrieved from the UniProt proteome database (https://www.uniprot.org/proteomes/UP000011083). Sequence alignment of CTSB and BRD4 against the *A. castellanii* proteome was performed using the Needleman–Wunsch algorithm with a BLOSUM62 scoring matrix [24]. Proteins with a sequence similarity ≥30% were considered potential nitroxoline targets within the *A. castellanii* proteome.

### 2.11. Homology modelling and molecular docking

Homology modelling and molecular docking were conducted using Schrödinger Suite 2022–3 (New York, NY, USA). Proteins and ligands were prepared using the Protein Preparation Wizard and LigPrep modules in the Schrödinger Suite with default settings. A receptor grid was generated using the Receptor Grid Generation module in Schrödinger Suite, and the ligand diameter midpoint boxes were defined as 10×10×10 Å^3^. Molecular docking was performed using the Ligand Docking module in the Schrödinger Suite with SP precision, a flexible ligand sampling strategy, and a van der Waals radii scaling factor of 0.5 for the receptor and ligand [25]. Subsequently, docking poses underwent Prime MM-GBSA computation, where residues within 6 Å of the docked ligands were relaxed using the minimise sampling method. Finally, the binding free energies (MM-GBSA dG Bind) and chemotype diversity were calculated by visually inspecting the optimised docking pose and considering the SP docking scores.

### 2.12. Monodansylcadaverine (MDC) autophagy staining assay in trophozoites

An MDC staining assay kit (Beyotime, C3018S, Nanjing, China) was used to detect autophagic vesicles in autophagic cells. All vacuoles in the trophozoites were visualised using MDC staining. Following the manufacturer’s instructions, trophozoites (1×10^5^) after a 12-h treatment were washed once with 1× PBS and incubated for 30 min at 26°C in 1 mL of assay buffer containing 1 μL of MDC, the tracer of acidic autophagic vesicles. After three washes with assay buffer, fluorescent MDC signal acquisition (λEx = 335 nm and λEm = 512 nm) was performed on a modular multimode microplate reader (BioTek Synergy H1).

### 2.13. Statistical analysis

All statistical analyses were performed using the GraphPad Prism 8 software (GraphPad Software, Version 8.0; San Diego, CA, USA). Comparisons between the control and treatment groups were performed using a one-way analysis of variance. Data are presented as means ± standard deviation (SD), derived from ≥3 independent experiments for each sample. Statistical significance was set at *p* < 0.05.

## 3. Results

3.1. Cell viability inhibition and morphology changes in *A. castellanii* after treatment with nitroxoline

To assess the in vitro efficacy of nitroxoline, we used the CellTiter-Glo® Luminescent cell viability assay. The determined IC50 value is presented in **Figure. 1A**, indicating that nitroxoline effectively inhibited *A. castellanii* trophozoites at 24 h with an IC50 ≤16.08±0.93 μM. **Figure 1B** depicts the time- and dose-dependent decreases in *A. castellanii* trophozoite viability in response to nitroxoline. Treatment with 20, 40, 60, and 80 μM of nitroxoline separately led to significant reductions in trophozoite viability to 32.14% ± 0.97%, 20.87% ± 0.22%, 17.29% ± 0.70%, and 9.85% ± 0.93% after 24 h (*p* < 0.05; **Fig. 1B**), respectively, compared with the 0.1% DMSO-treated group. Specifically, significant declines were observed at 20, 40, 60, and 80 μM of nitroxoline after 48 h (6.35% ± 0.48%, 2.42% ± 0.27%, 1.51% ± 0.10%, and 1.15% ± 0.04%) and 72 h (1.77% ± 0.26%, 0.55% ± 0.21%, 0.41% ± 0.02%, and 0.37% ± 0.06%) (*p* < 0.05; **Fig. 1B**). Data are presented as mean ± SD from triplicate experiments.

**Fig. 1.**
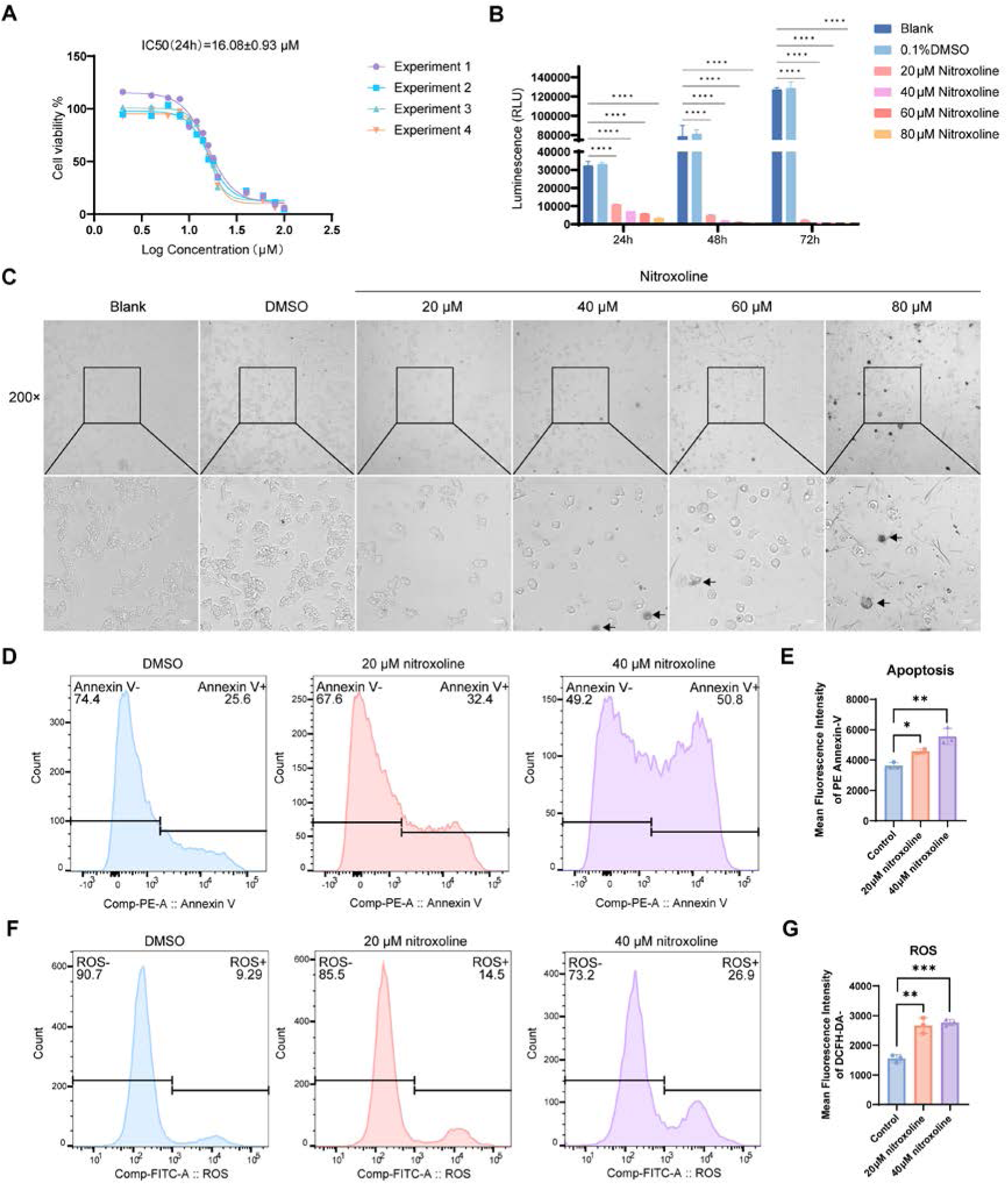
Suppressing *A. castellanii* trophozoites viability by nitroxoline. (A) IC50 values of nitroxoline-treated *A. castellanii* trophozoites determined by 24 h CellTiter-Glo assay. (B) Viability of trophozoites following 20–80 μM nitroxoline treatments for 24, 48, and 72 h determined using the CellTiter-Glo assay. Data are presented as means ± SDs from three experiments. Blank: without any treatments. DMSO-treated trophozoites (0.1%) were used as the negative control (\*\*\*\**p* < 0.0001). (C) Effects on *A. castellanii* trophozoites incubated with negative control (0.1%DMSO) and different concentration of nitroxoline for 24 h monitored under an inverted microscope (magnification, ×200). Scale bar: 20 μm. The arrows denote the rounded or lysed trophozoites penetrated with nitroxoline. (D) Trophozoites were treated with the indicated compound concentrations for 24 h. Apoptosis rate was measured using PE Annexin-V flow cytometry. (E) Mean fluorescence intensity of PE Annexin-V in each group. Data are presented as means ± SDs of three experiments (\**p* < 0.05, \*\**p* < 0.01). (F) Reactive oxygen specie level was measured using DCFH-DA flow cytometry. (G) Mean flunorescence intensity of DCFH-DA in each group. Data are represented as means ± SDs of three separate experiments (\*\**p* < 0.01, \*\*\**p* < 0.001). SDs, Standard deviations

Observation using an inverted microscope showed that the shape of the trophozoites was severely disrupted, and the boundary was blurred with increasing drug concentrations after 24 h (**Fig. 1C**). Trophozoites progressively became rounded and elliptical with spinous protrusions, and intracellular vacuoles disappeared and clustered together after 48 and 72 h of treatment (**Fig. S1**). In severely damaged trophozoites, broken holes on the surface of the cell membrane were observed; therefore, nitroxoline could penetrate and dye them (**Fig. 1C)**.

To investigate whether nitroxoline-induced inhibition of trophozoite viability was linked to apoptosis, we evaluated phosphatidylserine levels on apoptotic trophozoite membranes using Annexin V apoptosis detection. Flow cytometry analysis indicated that, compared with the control group, nitroxoline treatment significantly promoted cell apoptosis (**Fig. 1D to E**). Excessive accumulation of ROS can break cellular homeostasis, leading to oxidative stress [26]. ROS determined using DCFH-DA increased in the nitroxoline-treated groups compared with the normal-cultured group (**Fig. 1F to G)**. Nitroxoline treatment significantly decreased the MMP, indicating mitochondrial dysfunction (**Fig. S2**).

Collectively, these in vitro experiments demonstrate the inhibitory effect of nitroxoline on *A. castellanii* trophozoite growth.

### 3.2. General property and GO term enrichment analysis of transcriptome sequencing

Three parallel samples, separately from normal-cultured, DMSO-treated, 20 μM nitroxoline-treated, and 40 μM nitroxoline-treated groups after 24 h, respectively, were successfully subjected to RNA-seq (**Fig. S3A**). A principal component analysis plot further revealed distinctions among the four trophozoite groups (**Fig. S3B**). The expression profiles of the normal culture and DMSO-treated groups showed minimal differences. The gene expression patterns from the 20 μM nitroxoline-treated and 40 μM nitroxoline-treated groups were discrete compared with the other groups (**Fig. S3C)**, suggesting high variability of the expression profiles in nitroxoline-treated groups and indicating that many of the key variations in the genes in the expression profiles probably determined the drug efficacy to trophozoites. Volcano plots were also shown in which the blue and purple regions represent upregulated and downregulated genes with significant changes in differential abundance (log2FC < −1 or log2FC > 1; p < 0.05 based on Student’s t-test with false discovery rate correction) **(Fig. S3D to E)**.

To categorise the DEGs, GO term enrichment analysis was performed, which included three fundamental groups: biological processes (BP), cellular components (CC), and molecular functions (MF). BP analysis in the 20 μM nitroxoline-treated group indicated that regulated DEGs were predominantly involved in the cell wall macromolecule catabolic process, cell wall polysaccharide catabolic process, xylan catabolic process, cell fate specification, small molecule biosynthetic process, and L-glutamate biosynthetic process (**Fig. 2A**). Meanwhile, the significantly regulated DEGs in the CC category (**Fig. 2A**) were associated with components such as the intrinsic component of the membrane, integral component of the membrane, extracellular region, porous membrane, and subrhabdomeral cisterna. MF analysis (**Fig. 2A**) of 20 μM nitroxoline-treated DEGs revealed enrichment in oxidoreductase activity, FMN binding, iron ion binding, triglyceride lipase activity, and hydroxyacyl glutathione hydrolase activity. Nevertheless, in the 40 μM nitroxoline-treated group, BP analysis (**Fig. 2B**) revealed significant alterations in a multitude of genes related to cellular response to DNA damage stimulus, DNA repair, double-strand break repair via homologous recombination, recombination repair, and double-strand break repair. In the CC category **(Fig. 2B**), components such as spindles, condensed chromosomes, outer membranes, mitochondrial envelope, and smc5-smc6 complex were distinctly clustered. In the MF analysis, DEGs were enriched in ATP hydrolysis activity, ATP-dependent activity, FMN binding, 3-5 DNA helicase activity, and DNA binding.

**Fig. 2.**
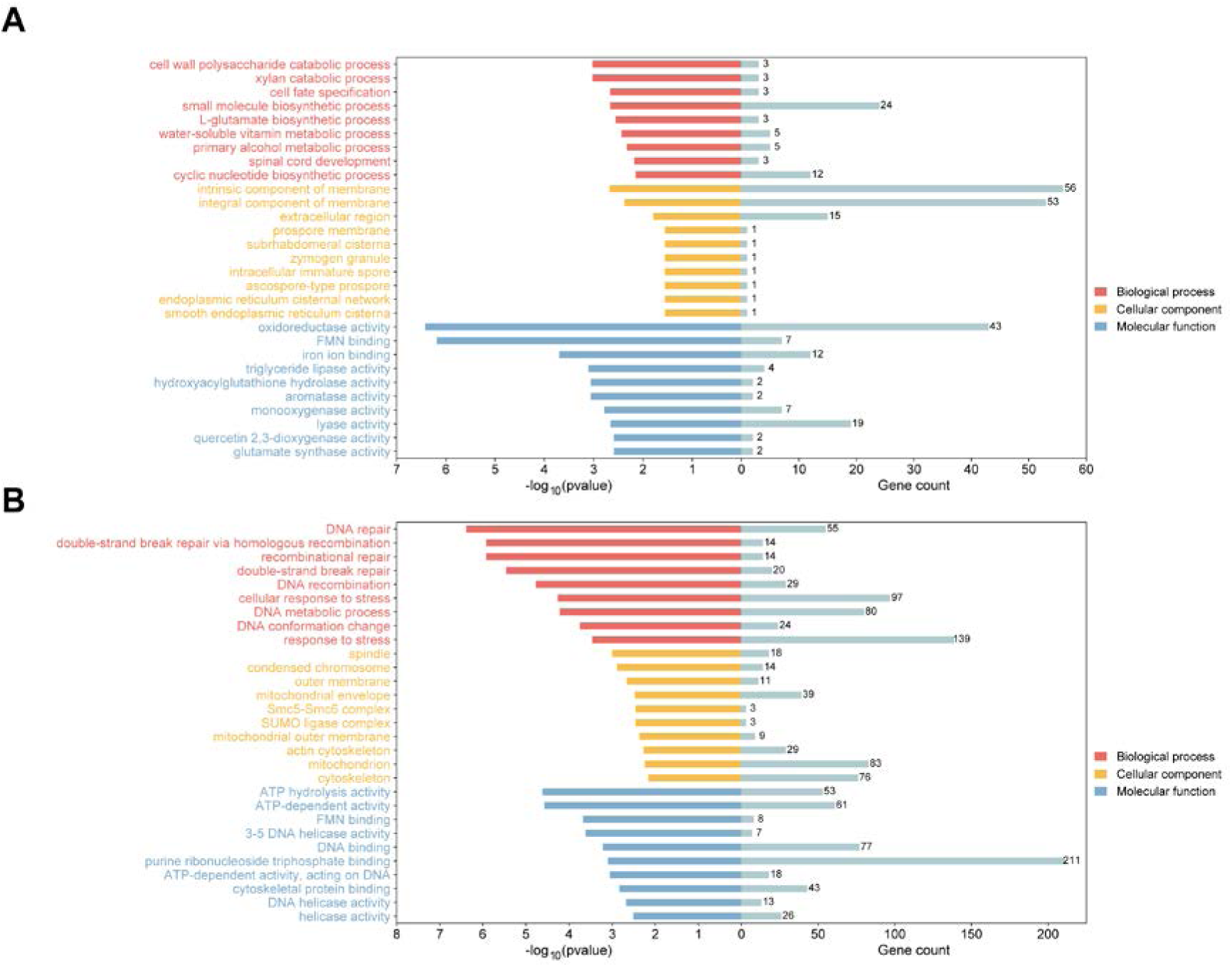
GO analysis to characterise functional pathways in *A. castellanii* trophozoites affected by nitroxoline. (A, B) Bar plot of top GO terms for significantly changed genes in 20 (A) and 40 μM nitroxoline-treated trophozoites (B). Red: biological process (BP). Yellow: cellular component (CC). Blue: molecular function (MF).

### 3.3. GO interaction network and KEGG pathway analysis

As depicted in the GO enrichment network, iron ion binding (GO:0005506) and oxidoreductase activity (GO:0016491) were enriched in the 20 μM nitroxoline-treated group (**Fig. 3A**), and cellular response to DNA damage stimulus (GO:0006974) was the top cluster of enrichment in the 40 μM nitroxoline-treated group (**Fig. 3B**). Pathway enrichment analysis based on KEGG terms revealed significant enrichment of regulated genes in hypertrophic cardiomyopathy; metabolic pathways; pyruvate metabolism; fatty acid biosynthesis; nitrogen metabolism; citrate cycle; alanine, aspartate, and glutamate; steroid degradation; C5-branched dibasic acid metabolism; and circadian rhythm metabolism in the 20 μM nitroxoline-treated group (**Fig. 3C**). Nonetheless, the genes in the 40 μM nitroxoline-treated group were significantly enriched in homologous recombination, fanconi anaemia pathway, base excision repair, biosynthesis of secondary metabolites, non-homologous end-joining, glycosphingolipid biosynthesis-globo series, plant hormone signal transduction, renin-angiotensin system, glycolysis/gluconeogenesis, and vibrio cholerae infection (**Fig. 3D**). Pathway enrichment analysis based on the GO interaction network and KEGG terms revealed significant enrichment of altered genes in various metabolic and DNA damage response (DDR) pathways. This transcriptomic response discrepancy to nitroxoline suggests that trophozoites adopt diverse regulatory mechanisms to counter external stress.

**Fig. 3.**
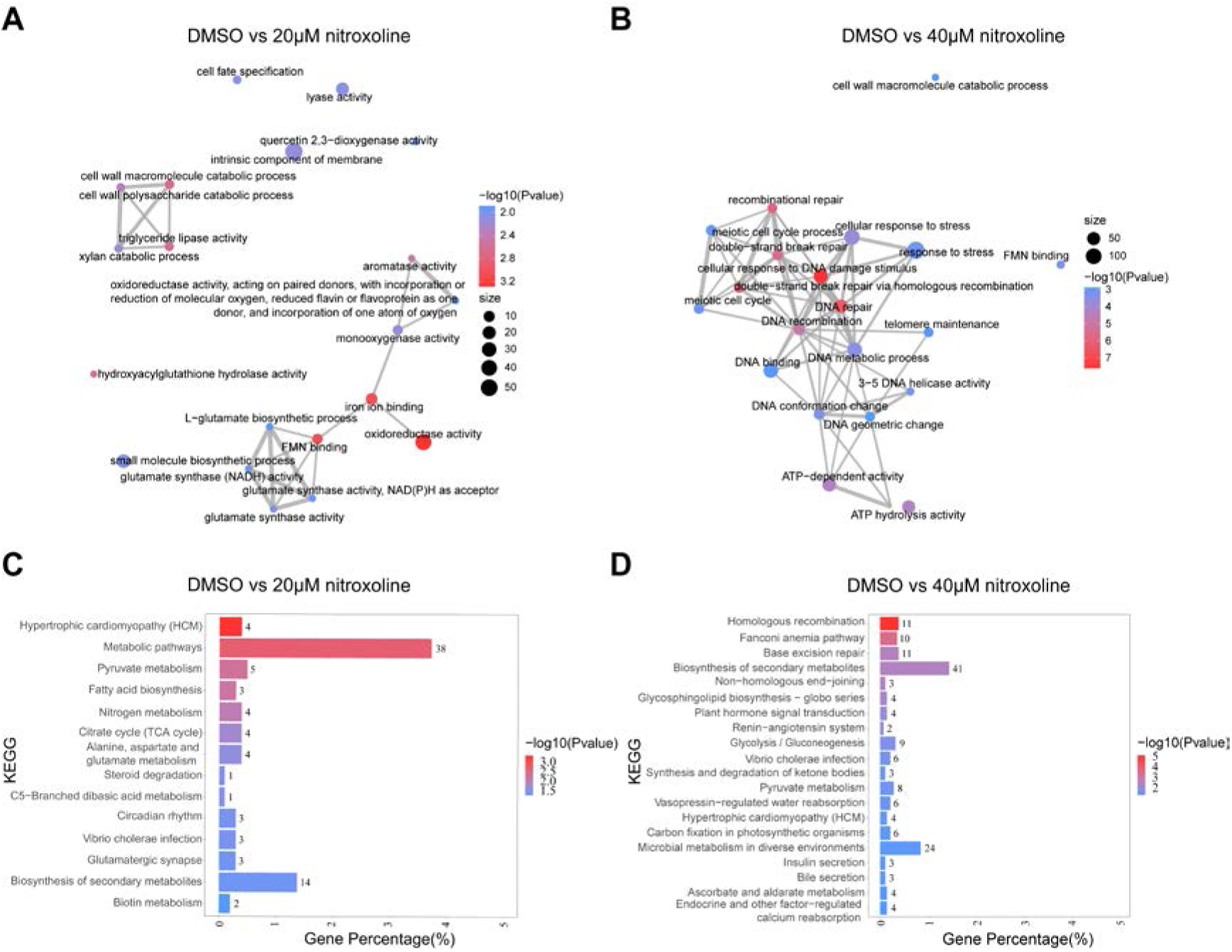
Network diagrams and KEGG analysis in *A. castellanii* trophozoites affected by nitroxoline. (A, B) Network diagrams depict the top 20 Gene Ontology terms interactions, grouped by pathways. Line thickness corresponds to the number of shared genes between two pathways. The ratio of overlapping genes to unique genes is ≥20%. The colour and size of the circles indicate the value of significantly regulated genes among each cluster. (C, D) Bar plot of top enriched KEGG terms for significantly changed genes in 20 and 40 μM nitroxoline-treated trophozoites. Statistical significance and gene percentage (number of genes in transcriptomic data versus all genes annotated with the KEGG term) are shown. KEGG, Kyoto Encyclopedia of Genes and Genomes

### 3.4. Nitroxoline alters the mRNA expression of trophozoites

The volcano plot showed that several DNA damage repair genes were upregulated after 24 h of treatment with nitroxoline (**Fig. 4A to B**), including *TOP2*, *PARG*, *BRCA2*, *FEN1*, and *RAD51*. In addition, several other DNA damage repair genes with significant upregulation following a 40 μM nitroxoline treatment of trophozoites were also observed (Fig. 4B), including *ATR, ATM, RAD50*, *RAD51 homolog*, *RAD54*, *RAD52*, *APE1*, *HDAC1*, *PARP149970* (UniProt IDs: L8HC40), *PARP256450* (UniProt IDs: L8GH34), *PARP114300* (UniProt IDs: L8H6I0), and *PARP338640* (UniProt IDs: L8HHX0). Upon further analysis of down-regulated genes following 40 μM nitroxoline treatment, the following genes were altered in trophozoites: *SAHH*, *GNMT*, *CBS*, *Cys1a*, and *MAT*. Genes (UniProt IDs: L8GGB2, L8GKJ9) belonging to the cytochrome p450 superfamily were upregulated in 20 μM and 40 μM nitroxoline treatments.

**Fig. 4.**
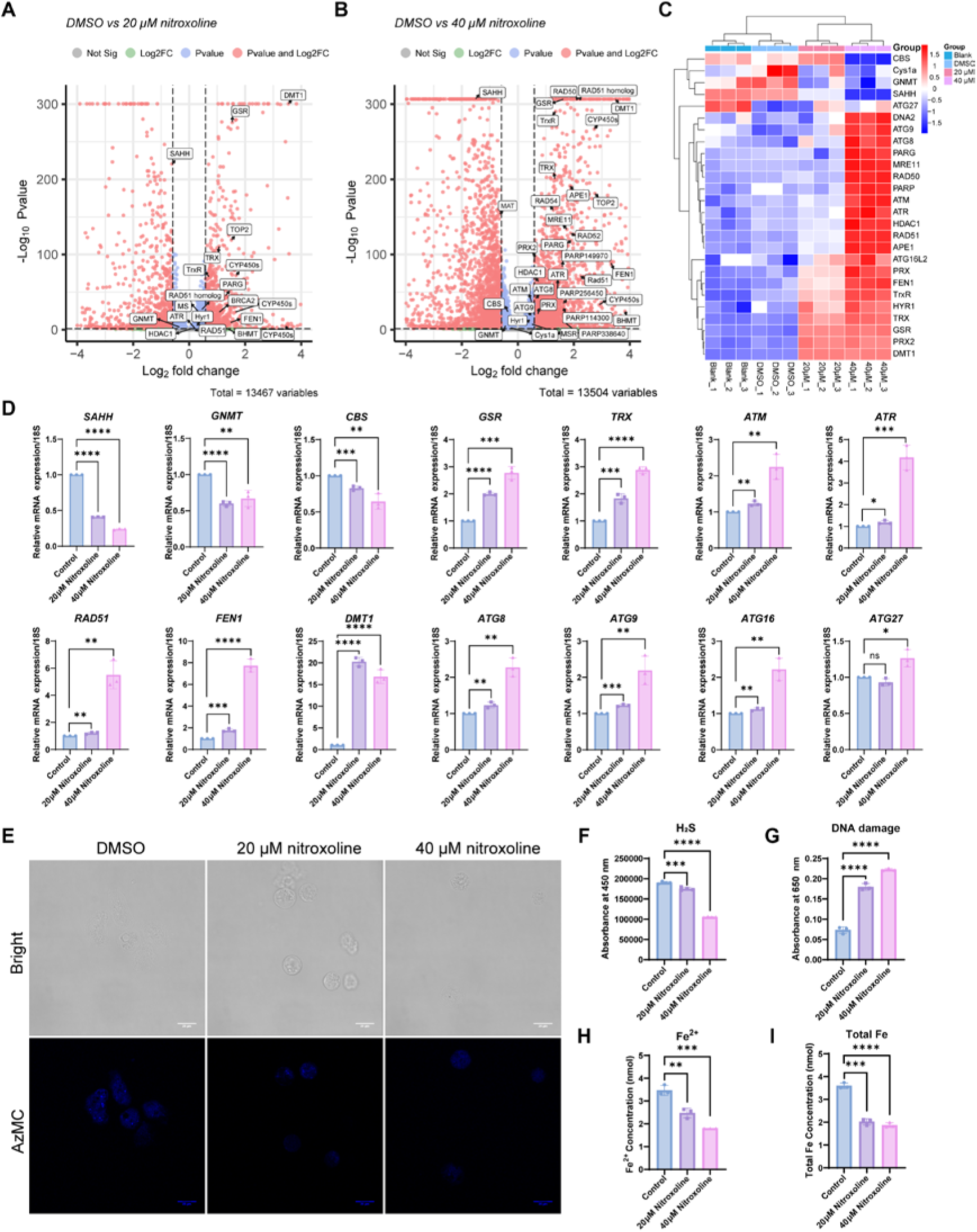
Transcript profiles in trophozoites treated with nitroxoline. (A, B) Enhanced volcano plot of the DEGs in trophozoites under 20 and 40 μM nitroxoline treatments, respectively. The metabolic pathways, DNA damage repair pathway and mitochondrial function related genes significant (|Fold change| >=1.5, Pvalue ≤0.05) are annotated. (C) Heat map of DEGs in nitroxoline-treated groups compared with DMSO-treated and normal-cultured groups (Blank). Increased and decreased abundances, relative to the control, are shown in red and blue, respectively. (D) Relative mRNA expression of *SAHH*, *GNMT*, *CBS*, *GSR*, *TRX*, *ATM*, *ATR*, *RAD51*, *FEN1*, *DMT1*, *ATG8*, *ATG9*, *ATG16*, and *ATG27* under 20 and 40 μM nitroxoline treatments for 24 h in *A. castellanii* trophozoites. Gene expression was normalised to *18s* expression levels. Results represent means ± standard deviations of three independent experiments (\**p* < 0.05, \*\**p* < 0.01, \*\*\**p* < 0.001, \*\*\*\**p* < 0.0001). (E) Representative H_2_S live-cell fluorescence images after treatment of nitroxoline. Trophozoites were predyed with 100 μM AzMC for 1 h. Scale bars, 20 μm. (F) Endogenous levels of H_2_S were detected using AzMC fluorescence assays (λEx = 365 nm, λEm = 450 nm). (G) DNA damage was assessed using avidin-biotin fluorescence assays (OD= 650 nm) (H, I) Iron amount (Fe^2+^ and Fe^3+^) was monitored using fluorescence assays (OD= 593 nm). DEGs, Differentially expressed genes; OD, Optical density

Visualising the selected DEGs within these pathways using a heatmap (**Fig. 4C**) revealed distinct gene expression patterns. Nitroxoline can positively or negatively affect metabolic enzymes. Treatment with 40 μM nitroxoline impacted methionine and cysteine cycles by decreasing the expression of *GNMT*, *SAHH*, *CBS,* and *Cys1a* (**Fig. 4C)** and increasing the expression of *BHMT* (**Fig. S4A to B**) in *A. castellanii*. In addition to the rescue of cell proliferation upon cysteine deprivation, *GNMT* expression normalised cellular oxidative stress, as determined by a reduction in intracellular ROS levels, and the transsulfuration pathway is required to sustain glutathione levels and redox balance in vivo [27]. Therefore, the loss of *GNMT* and *CBS* may disturb cellular oxidative homeostasis. Nevertheless, nitroxoline significantly activated various kinases, such as *ATM* and *ATR*, which act as the primary transducers in the DDR signalling cascade (**Fig. 4C**). Many of the downstream genes involved in DNA damage, such as *PARP*, *PARG*, *MRE11*, *RAD50*, *RAD51,* and *APE1*, exhibited high expression levels after 40 μM nitroxoline treatment (**Fig. 4C**). As the major contributors to detecting DNA damage and maintaining genomic stability, *HDAC1* showed increased expression in the 40 μM nitroxoline-treated group. Consequently, several physiologically important antioxidant enzymes of the peroxiredoxin, thioredoxin, and glutathione systems, such as *PRX*, *PRX2*, *TRX*, *TrxR*, *GSR,* and *HYR1*, were significantly upregulated in the 20 and 40 μM nitroxoline-treated groups (**Fig. 4C**). *ATG8*, *ATG9,* and *ATG16L2*, the autophagy-related genes, represented high mRNA expression levels after 40 μM nitroxoline treatment (**Fig. 4C**).

To further verify the consistency of the transcriptome results, RT-PCR was performed. The expression of the vital metabolic pathway genes *SAHH*, *GNMT,* and *CBS* exhibited significant downregulation after 20 and 40 μM nitroxoline treatment. In contrast, a significant increase in the redox system genes *GSR* and *TRX* and the DNA damage repair-related genes *ATM*, *ATR*, *RAD51,* and *FEN1* (**Fig. 4D**) were revealed in the 20 and 40 μM nitroxoline-treated groups. The expression of the other DNA repair relative genes *APE1*, *DNA2*, *PARP*, *PARG*, *MRE11,* and *RAD50* (**Fig. S4C to H**) increased only after 40 μM nitroxoline treatment, consistent with the transcriptomics. Moreover, the number of apurinic/apyrimidinic sites in DNA increased in the nitroxoline-treated groups, indicating probable nitroxoline-induced DNA damage (**Fig. 4G**). *DMT1*, involved in iron acquisition, was also upregulated in the 20 and 40 μM nitroxoline-treated groups (**Fig. 4D**). The autophagy-related genes *ATG8*, *ATG9*, and *ATG16* were significantly upregulated after 24 h of exposure to 20 or 40 μM nitroxoline (**Fig. 4D**), while *ATG27* remained relatively changed only after 40 μM nitroxoline treatment. Compared with the control group, only trophozoites treated with 40 μM nitroxoline showed more accumulation of autophagic vacuoles (**Fig. S4I**), consistent with the expression of autophagy-related genes.

### 3.5. Nitroxoline reduces the H_2_S production of trophozoites

8-Hydroxyquinoline derivatives can decrease the activity of *CBS* [28]. *CBS* encodes a pyridoxal 5′-phosphate-dependent enzyme that catalyses the condensation of homocysteine and serine to form cystathionine, which is a branch of the transsulfuration pathway. The transsulfuration pathway is the primary route for the biosynthesis of the major cellular antioxidants cysteine and glutathione, and cystathionine and its downstream partner CBS produce the gaseous transmitter H_2_S, another ROS scavenger, as a byproduct of their enzymatic activity [29]. Since CBS is also involved in H_2_S production, we measured its levels using an AzMC fluorogenic probe [30]. To investigate the effects of nitroxoline on trophozoites, H_2_S fluorescence imaging was performed using an AzMC probe (**Fig. 4E**). A 24-h treatment with 20 μM or 40 μM of nitroxoline significantly decreased H_2_S production levels by 7.90%

± 2.82% and 44.54% ± 0.82%, respectively, compared with the control group (**Fig. 4F**). Notably, nitroxoline reduced the level of endogenous H_2_S in a dose-dependent manner. Nitroxoline reduced H_2_S production in cellular lysates, ruling out a possible effect on regulating CBS expression levels or protein stability, which is similar to the above results showing that the mRNA expression level of CBS was significantly decreased after nitroxoline treatment for 24 h (**Fig. 4D**).

### 3.6. Nitroxoline impacts iron uptake transcripts in trophozoites

In a previous study [31], postulated that nitroxoline induces rapid iron starvation in the biofilms of several important pathogens. We were curious whether nitroxoline could affect iron uptake in *A. castellanii* trophozoites. Full transcriptomic analysis of nitroxoline-treated trophozoites revealed that the iron ion-binding pathways (**Fig. 3A**) were significantly affected. Nitroxoline treatment increased the transcription of genes involved in iron acquisition, including divalent metal transporter 1 (*DMT1)* (**Fig. 4C to D**). Bioinformatic analysis showed that *A. castellanii* genomes encode one gene product with homology to *Plasmodium falciparum* PfDMT1, *Arabidopsis thaliana* NRAMP2, and *Homo sapiens* DMT1 (NRAMP2). The DMT1 homologue was ACA1_225890 in *A. castellanii* (AcDMT1), which shared 58.31% amino acid sequence identity with *A. thaliana* NRAMP2, 56.67% with *H. sapiens* DMT1, and 28.38% with *P. falciparum* PfDMT1. DMT1 is essential for maintaining iron homeostasis and enables Fe^2+^ and Mn^2+^ ion entry into the mitochondria. Therefore, DMT1 promotes mitochondrial heme synthesis, iron-sulphur cluster biogenesis, and antioxidant defence [32]. The intracellular iron levels in nitroxoline-treated trophozoites were also confirmed (**Fig. 4H to I**). As depicted above, nitroxoline, especially at 40 μM, significantly decreased intracellular Fe^2+^ and total Fe amounts in trophozoites compared with the control group. These results establish a structural basis for the pharmacological intervention of nitroxoline on trophozoites.

### 3.7. Searching potential targets of nitroxoline in *A. castellanii* proteome

By searching ligand ’Nitroxoline’ in the RCSB PDB database, two entries (PDB codes: 3AI8 and 5Y1Y) were found with nitroxoline as a structural ligand. The first entry was human Cathepsin B (gene: CTSB, UniProt ID: P07858), which belongs to a family of lysosomal cysteine proteases known as cysteine cathepsins and plays an important role in intracellular proteolysis. The latter belongs to the human BET family of BRD4 (UniProt ID: O60885), a chromatin reader protein that recognises and binds to acetylated histones and plays a key role in transmitting epigenetic memory across cell divisions and transcription regulation. To identify possible *A. castellanii* proteins that are homologous to the known targets of nitroxoline (Cathepsin B and BRD4), we first downloaded the complete proteomic data of *A. castellanii* from the UniProt Proteomes database with 14,939 entries. The Needleman–Wunsch algorithm was used for sequence alignment using the BLOSUM62 scoring matrix. Cathepsin B and BRD4 sequences were globally aligned with all sequences in the *A. castellanii* proteome, followed by calculating sequence similarity. A sequence similarity cutoff (≥30%) was applied to isolate 639 homology proteins in *A. castellanii*. In addition, by comparing the GO annotation data of 639 homologous proteins with the GO term enrichment analysis outcomes illustrated in **Fig. 2A to C**, 69 *A. castellanii* proteins overlapped to potentially bind nitroxoline and associate with the DEGs caused by nitroxoline treatment. Homology modelling and molecular docking were performed for these 69 proteins using the Schrödinger Suite 2022–3, allowing for conformational flexibility of the targets and nitroxoline. As a positive control, we redocked the complex structure of cathepsin B with nitroxoline (PDB ID: 3AI8), obtaining a docking scoring of -4.30 and binding free energy (MMGBSA dG Bind) of -18.14 (**Fig. 5B**, **Table 1)**. After analysing the docking scores and binding poses, 18 proteins were identified as potential *A. castellanii* targets of nitroxoline (**Fig. 5**, **Table 1**).

**Fig. 5.**
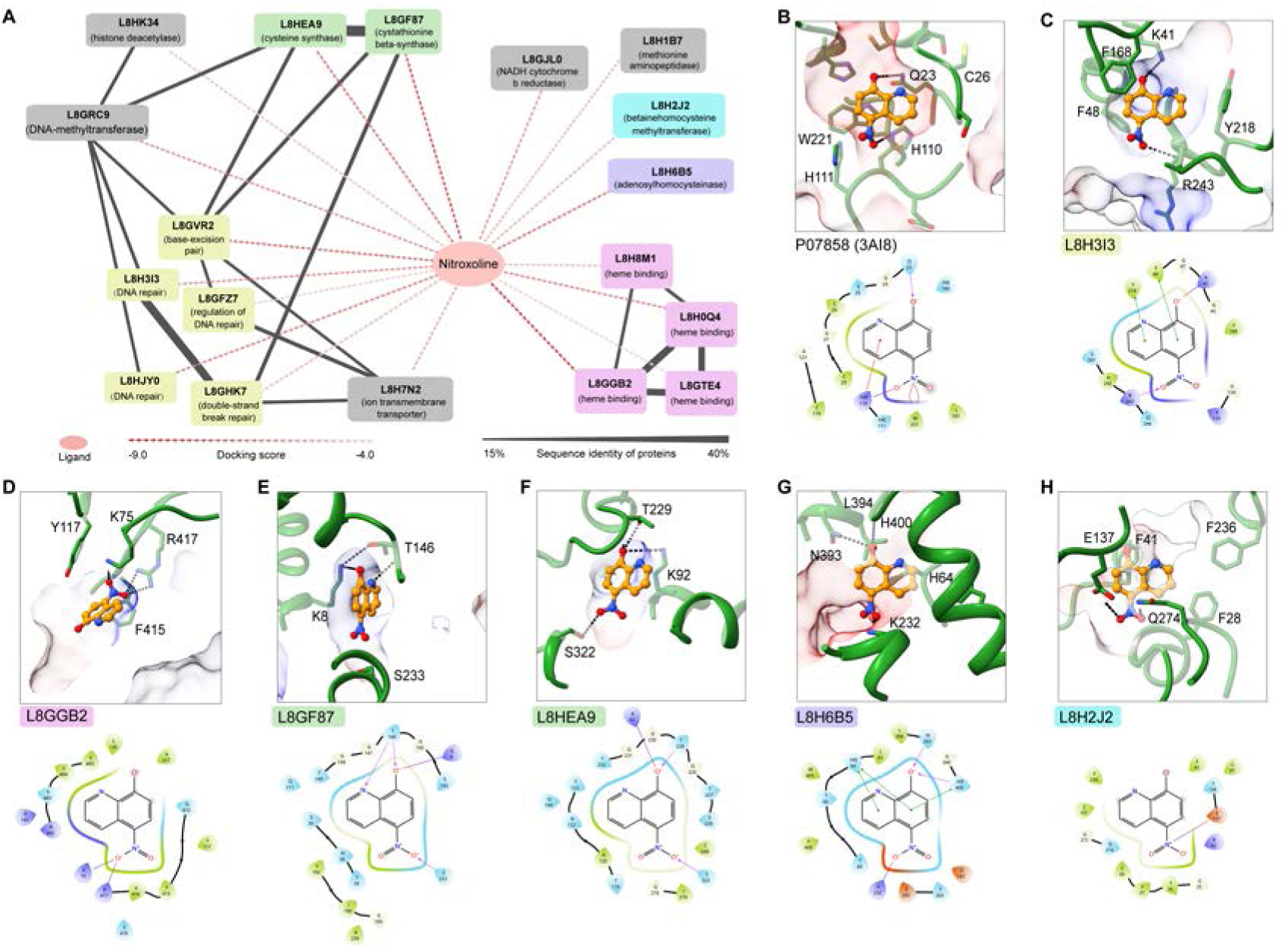
Protein similarity network and docking poses of nitroxoline in the potential *A. castellanii* targets. (A) Interaction network of nitroxoline with potential *A. castellanii* targets. (B–H) The surface representation and two-dimensional diagrams illustrate ligand–protein interactions within the predicted nitroxoline-binding pockets of potential targets, including positive control (B), DNA repair protein RAD51 (C), cytochrome p450 (D), cystathionine beta-synthase (E), cysteine synthase (F), adenosylhomocysteinase (G), and betainehomocysteine methyltransferase (H) The colour coding transitions from dodger blue, representing the most hydrophilic regions, to orange to red, indicating the most hydrophobic regions, within the surface representation. Key residues implicated in recognising nitroxoline are depicted as sticks. Polar interactions are denoted by black dashed lines. The amino acids of potential target residues are highlighted to indicate positive charge (blue), negative charge (red), polar (cyan), and hydrophobicity (green).

**Table 1.** List of the potential nitroxoline targets in *A. castellanii* that also associated with the Gene Ontology analysed functional pathways in *A. castellanii* trophozoites affected by nitroxoline.

| Group | Targets | Docking Score | MMGBSA dG Bind (kcal/mol) | Protein Names |
| --- | --- | --- | --- | --- |
| Proteins associated with DNA repair | **L8H3I3** | **-6.28** | **-50.86** | **DNA repair protein RAD51 homolog** |
|  | L8HJY0 | -6.52 | -25.29 | DNA repair protein |
|  | L8GFZ7 | -4.82 | -12.16 | Tyrosyl-tRNA synthetase |
|  | L8GVR2 | -7.16 | -1.33 | Flap endonuclease |
|  | L8GHK7 | -5.12 | 0.28 | Rad5111 protein |
| Cytochrome p450 family | L8H8M1 | -5.21 | -39.31 | Cytochrome P450 monooxygenase, putative |
|  | **L8GGB2** | **-8.47** | **-20.10** | **Cytochrome p450 superfamily protein** |
|  | L8GTE4 | -4.66 | -6.48 | Cytochrome P450, putative |
|  | L8H0Q4 | -6.48 | 8.02 | Cytochrome p450 superfamily protein |
| Cysteine synthase related proteins | **L8GF87** | **-8.08** | **-32.41** | **Cystathionine beta-synthase** |
|  | **L8HEA9** | **-7.04** | **-25.69** | **Cysteine synthase** |
| Others | L8GRC9 | -6.25 | -26.34 | DNA (cytosine-5)-methyltransferas |
|  | L8GJL0 | -6.20 | -16.38 | NADH cytochrome b reductase, putative |
|  | **L8H2J2** | **-5.42** | **-15.02** | **Betainehomocysteine methyltransferase** |
|  | **L8H6B5** | **-7.08** | **-10.49** | **Adenosylhomocysteinase** |
|  | L8H1B7 | -5.48 | -2.82 | Methionine aminopeptidase |
|  | L8H7N2 | -5.74 | -2.79 | Uncharacterized protein |
|  | L8HK34 | -5.14 | 4.25 | histone deacetylase |
| Control | **P07858 (3AI8)** | **-4.30** | **-18.14** | **Cathepsin B** |

The 18 potential targets were categorised into four groups. The first group included five proteins related to DNA repair that are crucial for protecting the genome from damage and mutations, thereby ensuring cell survival. The DNA repair protein RAD51 (UniProt ID: L8H3I3), which contains the Rad51 domain, had a good docking score of -6.28 and the best binding free energy (MMGBSA dG Bind) of - 50.86 kcal/mol, significantly better than that of the control. Nitroxoline was stabilised through multiple stacking interactions with F48 and Y218 and polar interactions with K41 and K243 (**Fig. 5C**). The second group comprised four proteins belonging to the cytochrome p450 family, which play essential roles in detoxifying and eliminating various compounds, the synthesis and breakdown of hormones, cholesterol synthesis, and vitamin D metabolism. L8GGB2, which contains the P450 domain, had the best docking score of -8.47 and a binding free energy of -20.10 kcal/mol. Nitroxoline formed salt bridges with K75 and R417 and engaged in extensive hydrophobic interactions with the surrounding residues (**Fig. 5D**). The third group comprised two proteins (UniProt IDs: L8GF87 and L8HEA9) associated with cysteine synthesis. By sharing similar binding pockets, they primarily interact with nitroxoline through polar interactions (via K8/K92, T146/T229, and S233/S322) and display high docking scores and binding free energies (-8.08 and -7.04; MMGBSA dG Bind: -32.41 and -25.69 kcal/mol, respectively) (**Fig. 5E to F**). Additionally, seven discrete proteins were classified as others (**Fig. 5A**, **Table 1)**. Among them, adenosylhomocysteinase (UniProt ID: L8H6B5) and betainehomocysteine methyltransferase (UniProt ID: L8H2J2) had relatively better docking scores with nitroxoline (-7.08 and -5.42), being stabilised through multiple stacking and polar interactions (**Figs. 5G to H**). Collectively, these bioinformatic analyses and molecular docking studies support the possibility of direct interactions between nitroxoline and *A. castellanii* proteins.

## 4. Discussion

We elucidated the effects of nitroxoline on *A. castellanii* trophozoites in biological activities and molecular mechanisms, as illustrated in the schematic (**Fig. 6**). Methods for evaluating RNA expression levels, such as microarrays and RNA-seq, have become valuable tools for assessing the dynamic properties of biological systems in a fast, broad, and reliable manner. In addition to technological aspects, developing bioinformatic analytical methods and tools has enormously aided our capacity to extract knowledge from omics technologies.

**Fig. 6.**
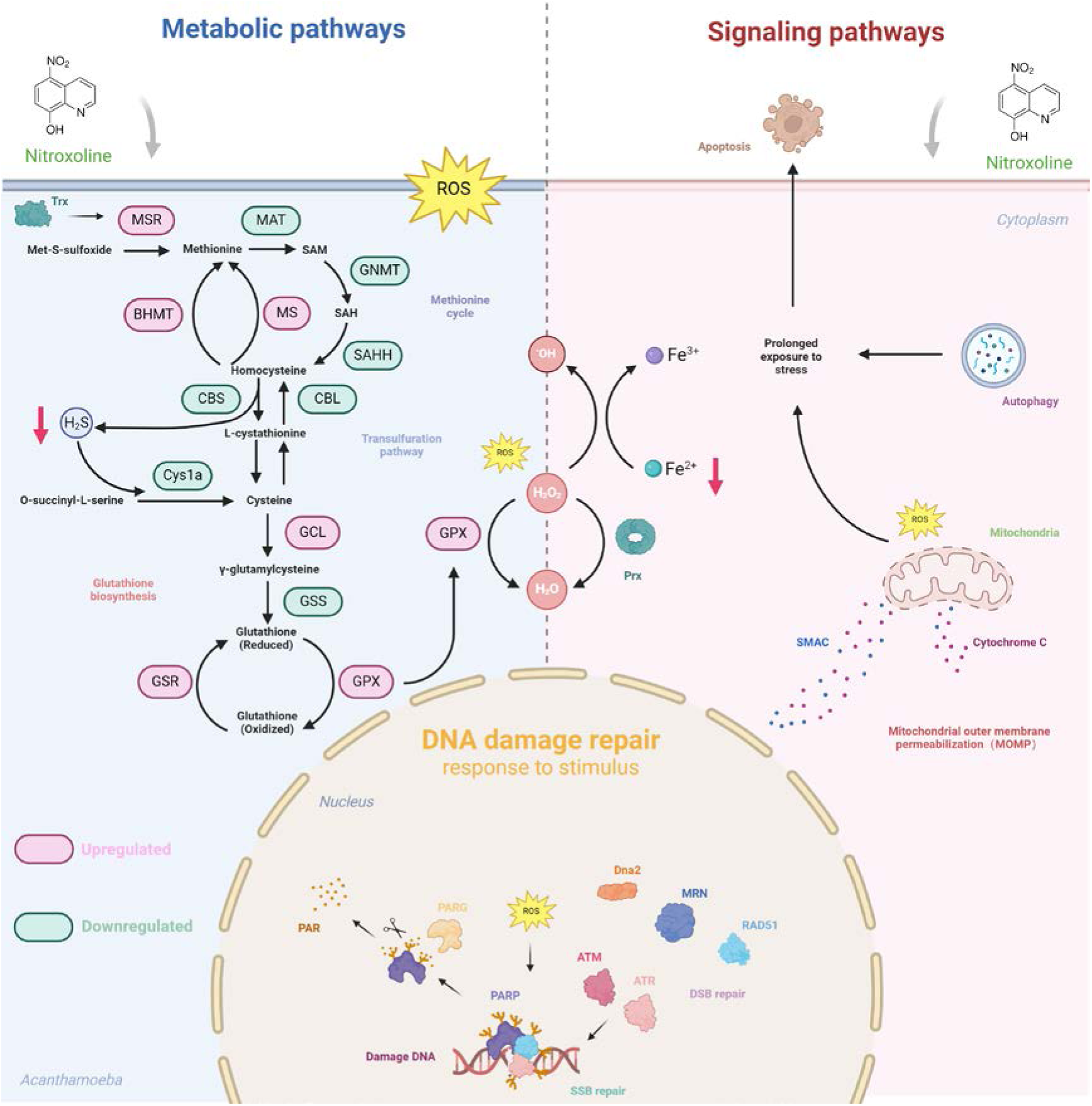
Schematic created with BioRender.com. MSR: Peptide-methionine (R)-S-oxide reductase. MAT: S-adenosylmethionine synthase. BHMT: Betainehomocysteine methyltransferase. MS: Methionine synthase. GNMT: DNA (cytosine-5)-methyltransferase. SAHH: S-adenosyl homocysteine hydrolase CBS: Cystathionine-β-synthase. CBL: Cystathionine-γ-ligase. Cys1a: Cysteine synthase. GCL: Glutamate-cysteine ligase. GSS: Glutathione synthetase. GSR: Glutathione reductase. GPX: Glutathione peroxidase. PARP: Poly [ADP-ribose] polymerase. PARG: Poly (ADP-ribose) glycohydrolase. ATM: Serine-protein kinase ATM family protein. ATR: Serine-protein kinase ATR family protein. Dna2: DNA replication factor Dna2. MRN: Meiotic recombination 11 homolog 1 (MRE11), DNA repair protein RAD50, and phosphopeptide-binding Nijmegen breakage syndrome protein 1 (NBS1). RAD51: DNA repair protein RAD51. Prx: Peroxiredoxin. Trx: Thioredoxin.

Reactive oxygen and nitrogen species are unavoidable byproducts of metabolic and energy transfer processes in oxidative life. These molecules play pivotal roles in regulating cellular processes. However, excess amounts of ROS or reactive nitrogen species cause oxidative and nitrosative stress, which may damage macromolecules. Free radicals generated in cells under oxidative stress can directly attack cellular membranes. Nitroxoline increased intracellular ROS levels in a dose-dependent manner. Consistent with this idea, nitroxoline treatment significantly increased *GPX* and *GSR* expression, indicating strong intracellular oxidation modulation. Additionally, following the momentum of big data research, *Trx*, *TrxR,* and *MsrA* were strongly upregulated after exposure to nitroxoline. Furthermore, *A. castellanii TrxR-S* reduces *Trx-1*, which reduces *Prx-1* and *MsrA,* suggesting that the synergistic interactions of these antioxidant enzymes are integral to the response to oxidative stress [33]. In a more recent study, mitochondrial fragmentation was accompanied by perturbations in energy production and elevated cellular ROS production [34]. In our study, nitroxoline significantly induced mitochondrial depolarisation in a dose-dependent manner.

Additional evidence for the role of nitroxoline in metabolic reprogramming comes from proteomic data in *E. coli,* highlighting the significant impact of this drug on metabolism [35]. When it comes to how nitroxoline impacted metabolic pathways, such as the methionine and cysteine cycles, the important clues come from the volcano plot of DEGs, which show the upregulated expression of *BHMT* and *MS* and downregulated expression of *GNMT*, *SAHH*, *CBS*, and *Cys1a*. The action of either *MS* or *BHMT* occurs during the resynthesis of methionine using Hcy. *GNMT* plays an important role in methionine clearance [36]. The result of these changes is a potential increase in methionine synthesis. Methionine is a major target of ROS, and its metabolism signals a state of nutrient deprivation [37,38]. A myraid of studies have shown that excess methionine also induces the transcription of ribosomal and rRNA genes and overall translation capacity [39]. Cysteine can be converted from o-succinyl-L-serine by *Cys1a* and is also a product of the transsulfuration pathway and precursor for the synthesis of tripeptide γ-glutamyl cysteinyl glycine (glutathione) [40,41]. Cysteine, homocysteine, glutathione, and cysteinyl glycine were the most abundant sulfhydryl groups. All these molecules are important in radical interactions as they are particularly susceptible to oxidation by ROS [42]. Subsequently, we found intriguing alternatives that nitroxoline could impact the expression of *CBS*, which is the first rate-limiting step in the transsulfuration pathway [27]. Additionally, we traced the effect of the reduction in *CBS* expression levels to its byproduct, H_2_S, and its role as a gaseous transmitter or ROS scavenger within the cell [37]. Following this hypothesis, H_2_S fluorescence imaging and qualitative analysis depicted nitroxoline substantially decreased the amount of H_2_S in *A. castellanii* trophozoites. H_2_S protects the cells from oxidative stress and modulates neuronal transmission, smooth muscle relaxation, insulin release, and inflammatory responses. In mammalian cells, endogenous H_2_S is produced by three enzymes: *CSE*, *CBS,* and *3-MST* [37]. However, *CSE* and *3-MST* are not annotated in the *A. castellanii* database. Since all these metabolic pathways and genomic integrity safeguards play crucial roles in maintaining cellular homeostasis and are critical for growth, they are acutely sensed and trigger signalling responses. Autophagy is required for maintaining cellular homeostasis. Autophagy occurs when trophozoites face drug stress and subsequently, experience a degree of amino acid starvation. Therefore, whether nitroxoline could contribute to autophagy is conceivable. Genetic analyses have identified that several autophagy-related genes (such as *ATG8*, *ATG9,* and *ATG16*) that execute and regulate autophagy were significantly upregulated. ATGs are organised into functional complexes that participate in each step of macroautophagy. Upregulating autophagy genes have also been observed in flies lacking *GNMT* [43]. An imbalance in HCY, glutathione, and cysteine metabolism can modulate autophagy [44–46]. In our study, nitroxoline significantly decreased the expression of *GNMT* and *CBS*, indicating a potential relationship between autophagy and metabolic pathways. Cellular oxidative stress is also key to autophagic response modulation [38]. Eventually, after prolonged exposure to nitroxoline, the trophozoites transitioned from autophagy to apoptosis.

Nitroxoline can cause DNA damage in trophozoites at high concentrations, which is consistent with the upregulated mRNA expression of an arsenal of enzymes capable of sensing replication stress and transducing information to influence cellular responses and neutralise the destructive effects of aberrant DNA structures. Transcriptomic and qPCR analyses showed the upregulation of *ATM*, *ATR*, *RAD51*, *RAD50*, *MRE11,* and *FEN1*. The DNA damage repair is a complex signal transduction pathway [47]. The first step in DSBs is recruiting repair factors at DNA break sites, which are initially recognised by the *MRE11*-*RAD50*-*NBS1* complex and promote ATM activation and DNA preparation for HR [48,49]. *RAD51*, a key protein that corrects errors in replication forks, is a canonical marker of DNA damage [50,51]. Volcano analysis also revealed several significant *PARPs*, such as *PARP* (UniProt IDs: L8HC40), *PARP* (UniProt IDs: L8GH34), *PAPR* (UniProt IDs: L8H6I0), and *PARP* (UniProt IDs: L8HHX0). *PARP* is a cellular stress sensor that is activated by oxidative, metabolic, and genotoxic stresses, such as single-strand break repair and double-strand DNA breaks, and in response, directs cells to specific fates according to the type and strength of the stress stimulus.

Homology modelling and molecular docking were performed to identify the potential targets of nitroxoline. Four main groups were categorised: five proteins related to DNA repair, four belonging to the cytochrome p450 family, two (UniProt IDs: L8GF87 and L8HEA9) associated with cysteine synthesis, and seven classified as others. The DNA repair protein RAD51 (UniProt IDs: L8H3I3), which contains the Rad51 domain, had a good docking score of -6.28 and the best binding free energy (MMGBSA dG Bind) of -50.86 kcal/mol. Cys1a (UniProt IDs: L8HEA9) and CBS (UniProt IDs: L8GF87) shared similar binding pockets, primarily interacted with nitroxoline through polar interactions (via K8/K92, T146/T229, and S233/S322), and displayed high docking scores and binding free energies. SAHH (UniProt ID: L8H6B5) and BHMT (UniProt ID: L8H2J2) had relatively better docking scores with nitroxoline (-7.08 and -5.42), being stabilised through multiple stacking and polar interactions. Collectively, these transcriptomic analyses and molecular docking studies supported the possibility of direct interactions between nitroxoline and *A. castellanii* proteins.

Previous studies suggested that nitroxoline exerts its antibacterial effects by chelating metal ions such as Fe^2+^, Mn^2+^, Mg^2+^, and Ca^2+^ [52,53]. Iron and copper are transition metal ions that can cleave hydroperoxides to form radicals that initiate chain reactions [42]. Our findings suggest that the intracellular iron levels (Fe^2+^ and Fe^3+^) of nitroxoline-treated trophozoites were significantly decreased, and the transcription of *DMT1* (UniProt IDs: L8H7N2), which is involved in iron acquisition, was increased. The *A. castellanii* genome encodes a gene with homology to *Plasmodium falciparum* PfDMT1, *Arabidopsis thaliana* NRAMP2, and *Homo sapiens* DMT1 (NRAMP2). The DMT1 homologue was AcDMT1, which shared 58.31% amino acid sequence identity with *A. thaliana* NRAMP2, 56.67% with *H. sapiens* DMT1, and 28.38% with *P. falciparum* PfDMT1. The observed increase in the expression of intracellular ferrous iron transporters suggests a potential increase in intracellular recycling or the liberation of stored iron, aiding in sustaining growth within these comparatively brief timeframes [41]. Furthermore, the regulatory mechanisms underlying the changes in iron metabolism remain unclear. Further studies are underway to test these hypotheses.

Putting these nascent studies together, an early mechanistic picture began to coalesce that nitroxoline could impact the viability of *A. castellanii* trophozoites in a dose- and time-dependent manner, with increased metabolic process disorders, intracellular ROS production, DNA damage, iron starvation, decreased endogenous H_2_S, decreased mitochondrial outer membrane permeabilisation, and the possibility of direct interactions between nitroxoline and *A. castellanii* proteins.

## Statements and declarations

### Author contributions

XC and LC designed the study. LC and WH performed the experimental works. QZ helped with experiments and provided technical support. LC wrote the original draft of the paper, and WH, MF, QZ and XC reviewed and edited the draft. Funding was obtained by XC and QZ.

### Data availability

The original contributions of this study are included in the article/supplementary material section.

### Ethics approval

Not applicable.

## Acknowledgments

We thank Shuhui Sun and Shanghai NewCore Biotechnology Co., Ltd. (https://www.bioinformatics.com.cn, last accessed on 10 Nov 2023) for technical assistance.

## Supplements

**Supplementary Fig. 1.**
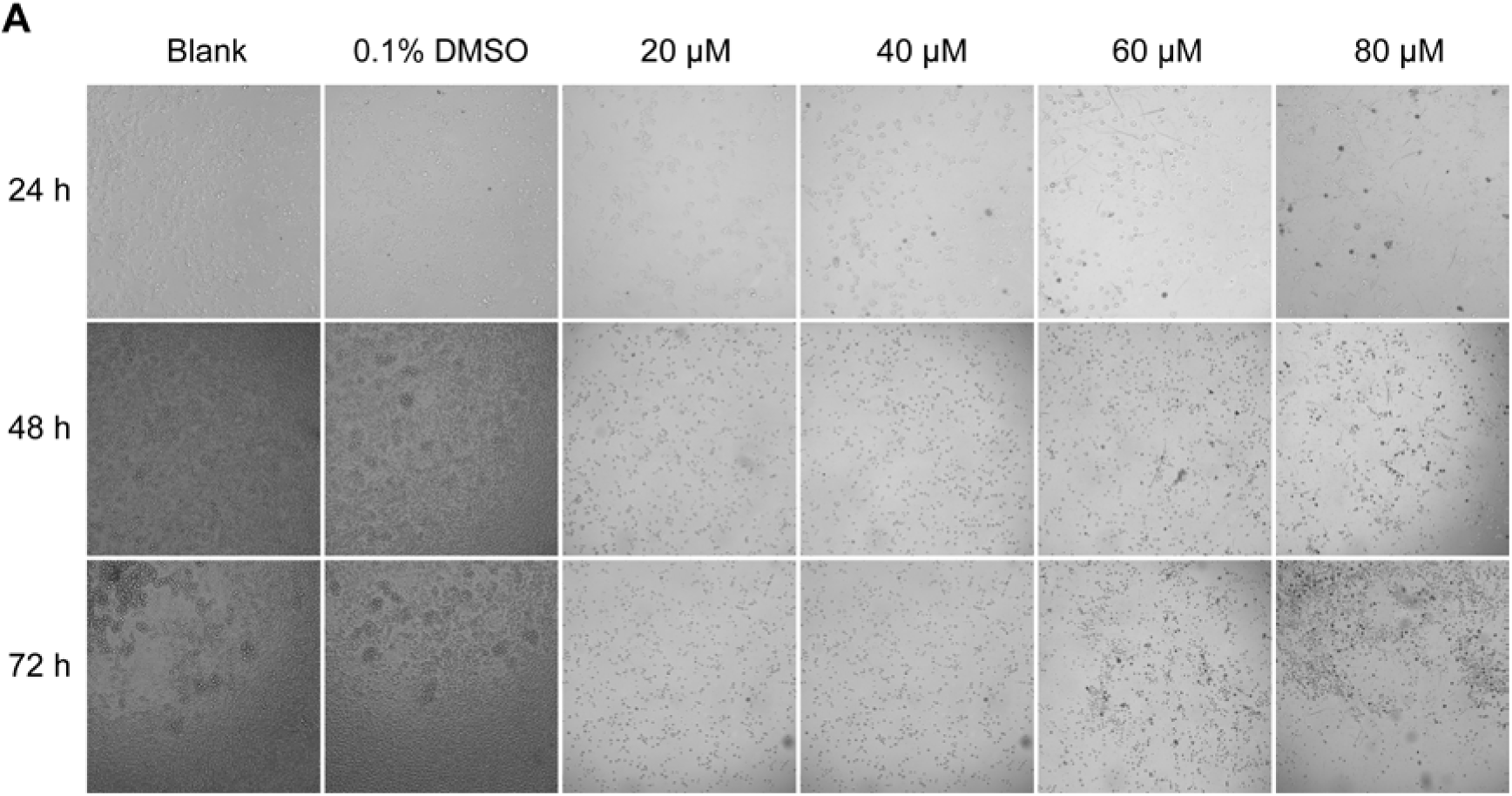
Effects on *A. castellanii* trophozoites incubated with negative control (0.1%DMSO) and different concentration of nitroxoline for 24, 48, and 72 h as monitored under an inverted microscope (magnification, ×200).

**Supplementary Fig. 2.**
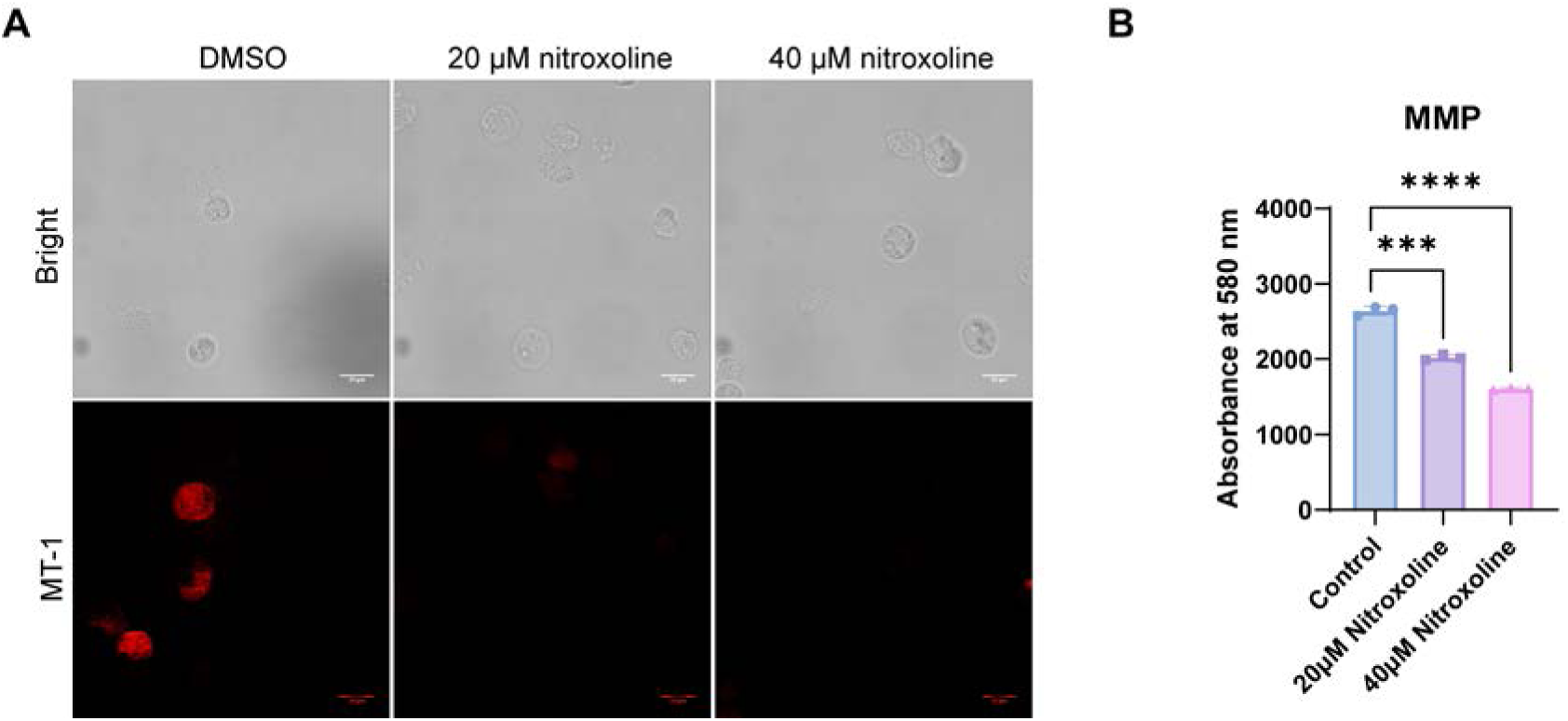
(A) Representative MT-1 fluorescence live-cell images after treatment of nitroxoline. Scale bars, 20 μm. (**B**) Mitochondrial membrane potential was evaluated using the MT-1 fluorescence assays (λEx = 488 nm, λEm = 580 nm).

**Supplementary Fig. 3.**
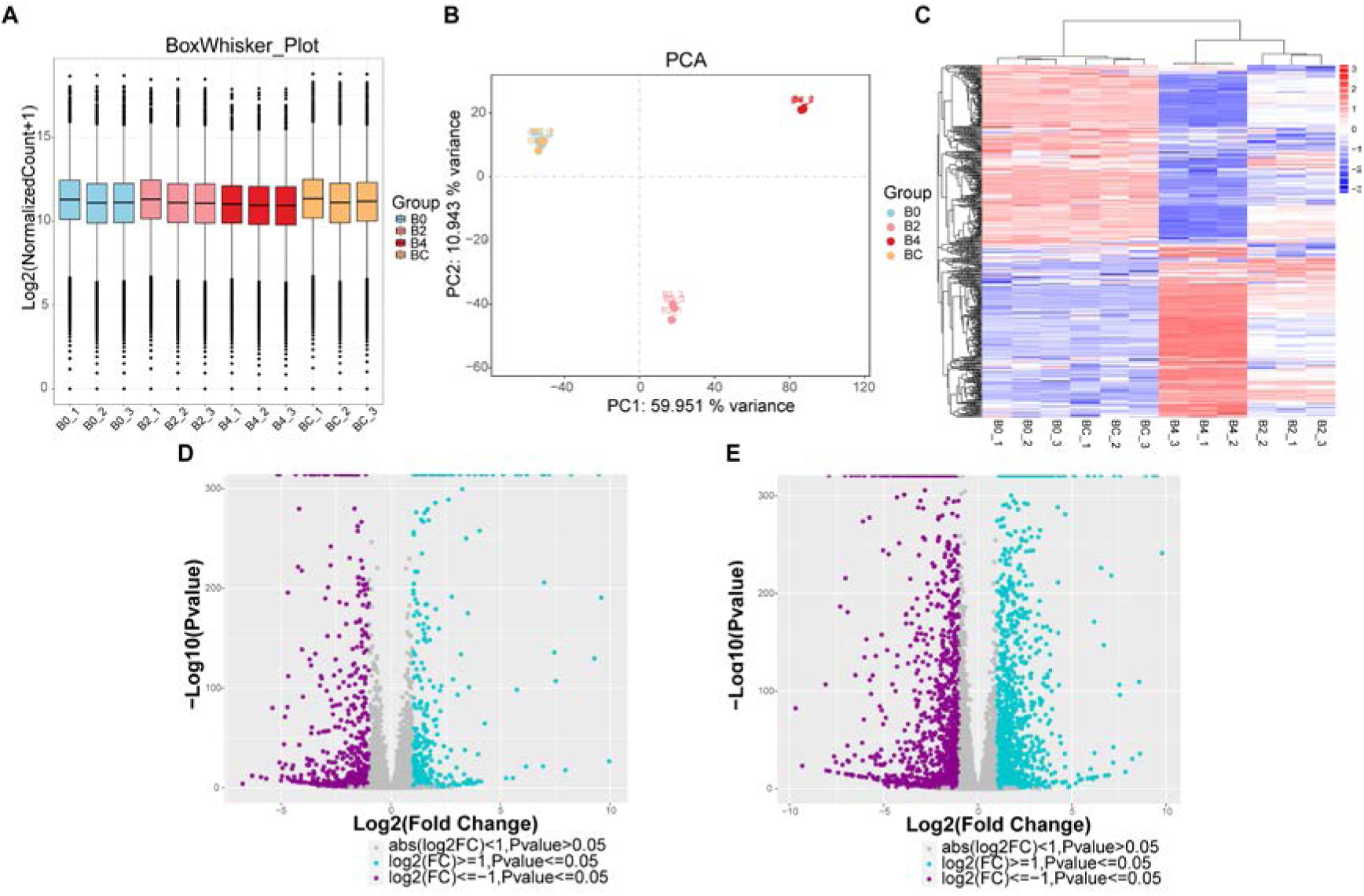
Quality assessment of transcriptome sequencing. (A) Average differentially expressed gene levels for four groups in RNA-seq analysis. B0: DMSO-treatment group, B2: 20 μM nitroxoline-treatment group, B4: 40 μM nitroxoline-treatment group, BC: normal cultured group. (B) Principal component analysis in the four groups was analysed using transcriptome sequencing. (C) Heat map of gene differential expression levels in 20 and 40 μM nitroxoline-treated trophozoites compared with control (DMSO-treated). Volcano plots of expression fold change in 20 μM nitroxoline-treated group versus DMSO-treated group (D) and 40 μM nitroxoline-treated group versus DMSO-treated group (E). (D, E) Upregulated genes are shown in blue, whereas downregulated genes are shown in purple.

**Supplementary Fig. 4.**
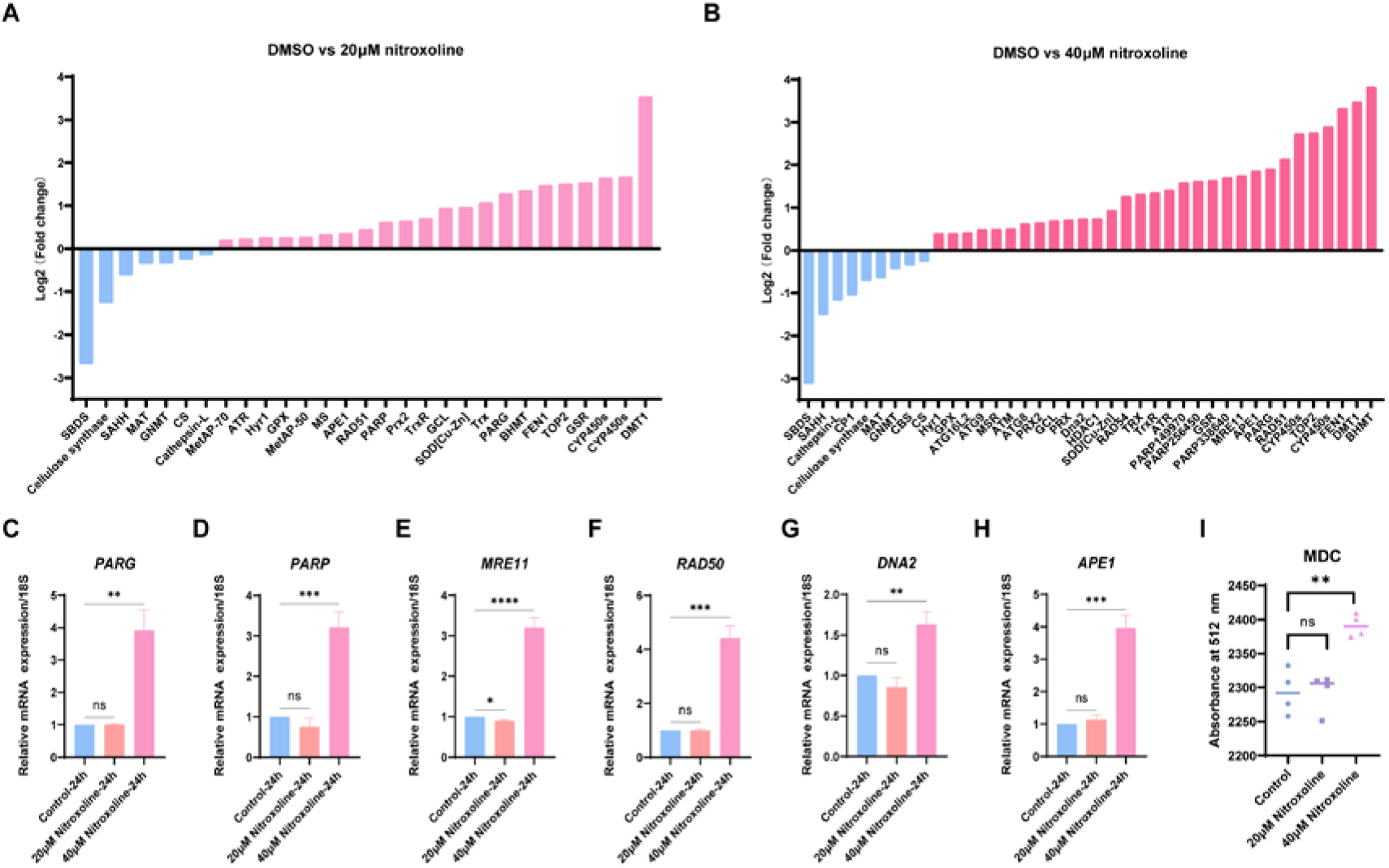
(A, B) Bar plot displaying the log2 (fold change) for the differentially expressed genes in trophozoites under 20 and 40 μM nitroxoline treatment, respectively. Each bar represents an *A. castellanii* gene. (C–H) Relative mRNA expression of *PARG*, *PARP*, *MRE11*, *RAD50*, *DNA2*, and *APE1* under 20 and 40 μM nitroxoline treatment for 24 h in *A. castellanii* trophozoites. (I) Fluorescent autophagy signal acquisition (λEx = 335 nm and λEm = 512 nm) was performed using MDC dying.

## References

[1] H. Zhang, X. Cheng, Various brain-eating amoebae: the protozoa, the pathogenesis, and the disease, Front Med 15 (2021) 842–866. 10.1007/s11684-021-0865-2.

[2] G.S. Visvesvara, H. Moura, F.L. Schuster, Pathogenic and opportunistic free-living amoebae: Acanthamoeba spp., Balamuthia mandrillaris, Naegleria fowleri, and Sappinia diploidea, FEMS Immunol Med Microbiol 50 (2007) 1–26. 10.1111/j.1574-695X.2007.00232.x.

[3] R. Siddiqui, N.A. Khan, Biology and pathogenesis of Acanthamoeba, Parasit Vectors 5 (2012) 6. 10.1186/1756-3305-5-6.

[4] M. Cabrera-Aguas, P. Khoo, S.L. Watson, Infectious keratitis: A review, Clin Exp Ophthalmol 50 (2022) 543–562. 10.1111/ceo.14113.

[5] J.Y. Niederkorn, The biology of Acanthamoeba keratitis, Exp Eye Res 202 (2021) 108365. 10.1016/j.exer.2020.108365.

[6] S.K. Kalra, P. Sharma, K. Shyam, N. Tejan, U. Ghoshal, Acanthamoeba and its pathogenic role in granulomatous amebic encephalitis, Exp Parasitol 208 (2020) 107788. 10.1016/j.exppara.2019.107788.

[7] Y.-J. Wang, C.-H. Chen, J.-W. Chen, W.-C. Lin, Commensals Serve as Natural Barriers to Mammalian Cells during Acanthamoeba castellanii Invasion, Microbiol Spectr 9 (2021) e0051221. 10.1128/Spectrum.00512-21.

[8] G.L. Damhorst, A. Watts, A. Hernandez-Romieu, N. Mel, M. Palmore, I.K.M. Ali, S.G. Neill, A. Kalapila, J.R. Cope, Acanthamoeba castellanii encephalitis in a patient with AIDS: a case report and literature review, Lancet Infect Dis 22 (2022) e59–e65. 10.1016/S1473-3099(20)30933-6.

[9] S. Bonini, A. Di Zazzo, G. Varacalli, M. Coassin, Acanthamoeba Keratitis: Perspectives for Patients, Curr Eye Res 46 (2021) 771–776. 10.1080/02713683.2020.1846753.

[10] S. Haapanen, H. Barker, F. Carta, C.T. Supuran, S. Parkkila, Novel Drug Screening Assay for Acanthamoeba castellanii and the Anti-Amoebic Effect of Carbonic Anhydrase Inhibitors, J Med Chem 67 (2024) 152–164. 10.1021/acs.jmedchem.3c01020.

[11] F. O’Grady, B. Smith, Neuromyopathy in the mouse produced by the antimicrobial agent nitroxoline, J Pathol Bacteriol 92 (1966) 43–48. 10.1002/path.1700920106.

[12] A.R. Joaquim, M.P. Gionbelli, G. Gosmann, A.M. Fuentefria, M.S. Lopes, S. Fernandes de Andrade, Novel Antimicrobial 8-Hydroxyquinoline-Based Agents: Current Development, Structure-Activity Relationships, and Perspectives, J Med Chem 64 (2021) 16349–16379. 10.1021/acs.jmedchem.1c01318.

[13] F. Fuchs, H. Hof, S. Hofmann, O. Kurzai, J.F. Meis, A. Hamprecht, Antifungal activity of nitroxoline against Candida auris isolates, Clin Microbiol Infect 27 (2021) 1697.e7-1697.e10. 10.1016/j.cmi.2021.06.035.

[14] F. Fuchs, F. Becerra-Aparicio, K. Xanthopoulou, H. Seifert, P.G. Higgins, In vitro activity of nitroxoline against carbapenem-resistant Acinetobacter baumannii isolated from the urinary tract, J Antimicrob Chemother 77 (2022) 1912–1915. 10.1093/jac/dkac123.

[15] F. Fuchs, A.M. Aldejohann, A.M. Hoffmann, G. Walther, O. Kurzai, A.G. Hamprecht, In Vitro Activity of Nitroxoline in Antifungal-Resistant Candida Species Isolated from the Urinary Tract, Antimicrob Agents Chemother 66 (2022) e0226521. 10.1128/aac.02265-21.

[16] A.M. Hoffmann, M. Wolke, J. Rybniker, G. Plum, F. Fuchs, Activity of the old antimicrobial nitroxoline against Mycobacterium abscessus complex isolates, J Glob Antimicrob Resist 33 (2023) 1–4. 10.1016/j.jgar.2023.02.010.

[17] D. Bojkova, N. Zöller, M. Tietgen, K. Steinhorst, M. Bechtel, T. Rothenburger, J.D. Kandler, J. Schneider, V.M. Corman, S. Ciesek, H.F. Rabenau, M.N. Wass, S. Kippenberger, S. Göttig, M. Michaelis, J. Cinatl, Repurposing of the antibiotic nitroxoline for the treatment of mpox, J Med Virol 95 (2023) e28652. 10.1002/jmv.28652.

[18] J.C. Haston, J.R. Cope, Amebic encephalitis and meningoencephalitis: an update on epidemiology, diagnostic methods, and treatment, CNS Infections 36 (2023).

[19] R.L. Rodríguez-Expósito, I. Sifaoui, M. Reyes-Batlle, F. Fuchs, P.L. Scheid, J.E. Piñero, R. Sutak, J. Lorenzo-Morales, Induction of Programmed Cell Death in Acanthamoeba culbertsoni by the Repurposed Compound Nitroxoline, Antioxidants (Basel) 12 (2023) 2081. 10.3390/antiox12122081.

[20] M.T. Laurie, C.V. White, H. Retallack, W. Wu, M.S. Moser, J.A. Sakanari, K. Ang, C. Wilson, M.R. Arkin, J.L. DeRisi, Functional Assessment of 2,177 U.S. and International Drugs Identifies the Quinoline Nitroxoline as a Potent Amoebicidal Agent against the Pathogen Balamuthia mandrillaris, mBio 9 (2018) e02051–18. 10.1128/mBio.02051-18.

[21] L. Chen, W. Han, W. Jing, M. Feng, Q. Zhou, X. Cheng, Novel anti-Acanthamoeba effects elicited by a repurposed poly (ADP-ribose) polymerase inhibitor AZ9482, Front Cell Infect Microbiol 14 (2024) 1414135. 10.3389/fcimb.2024.1414135.

[22] UniProt Consortium, UniProt: the Universal Protein Knowledgebase in 2023, Nucleic Acids Res 51 (2023) D523–D531. 10.1093/nar/gkac1052.

[23] S.K. Burley, C. Bhikadiya, C. Bi, S. Bittrich, H. Chao, L. Chen, P.A. Craig, G.V. Crichlow, K. Dalenberg, J.M. Duarte, S. Dutta, M. Fayazi, Z. Feng, J.W. Flatt, S. Ganesan, S. Ghosh, D.S. Goodsell, R.K. Green, V. Guranovic, J. Henry, B.P. Hudson, I. Khokhriakov, C.L. Lawson, Y. Liang, R. Lowe, E. Peisach, I. Persikova, D.W. Piehl, Y. Rose, A. Sali, J. Segura, M. Sekharan, C. Shao, B. Vallat, M. Voigt, B. Webb, J.D. Westbrook, S. Whetstone, J.Y. Young, A. Zalevsky, C. Zardecki, RCSB Protein Data Bank (RCSB.org): delivery of experimentally-determined PDB structures alongside one million computed structure models of proteins from artificial intelligence/machine learning, Nucleic Acids Res 51 (2023) D488–D508. 10.1093/nar/gkac1077.

[24] S.B. Needleman, C.D. Wunsch, A general method applicable to the search for similarities in the amino acid sequence of two proteins, J Mol Biol 48 (1970) 443–453. 10.1016/0022-2836(70)90057-4.

[25] R.A. Friesner, R.B. Murphy, M.P. Repasky, L.L. Frye, J.R. Greenwood, T.A. Halgren, P.C. Sanschagrin, D.T. Mainz, Extra precision glide: docking and scoring incorporating a model of hydrophobic enclosure for protein-ligand complexes, J Med Chem 49 (2006) 6177–6196. 10.1021/jm051256o.

[26] Z. Liu, W. Tang, J. Liu, Y. Han, Q. Yan, Y. Dong, X. Liu, D. Yang, G. Ma, H. Cao, A novel sprayable thermosensitive hydrogel coupled with zinc modified metformin promotes the healing of skin wound, Bioact Mater 20 (2023) 610–626. 10.1016/j.bioactmat.2022.06.008.

[27] J. Zhu, M. Berisa, S. Schwörer, W. Qin, J.R. Cross, C.B. Thompson, Transsulfuration Activity Can Support Cell Growth upon Extracellular Cysteine Limitation, Cell Metab 30 (2019) 865–876.e5. 10.1016/j.cmet.2019.09.009.

[28] P. Conan, A. Léon, M. Gourdel, C. Rollet, L. Chaïr, N. Caroff, N. Le Goux, C. Le Jossic-Corcos, M. Sinane, L. Gentile, L. Maillebouis, N. Loaëc, J. Martin, M. Vilaire, L. Corcos, O. Mignen, M. Croyal, C. Voisset, F. Bihel, G. Friocourt, Identification of 8-Hydroxyquinoline Derivatives That Decrease Cystathionine Beta Synthase (CBS) Activity, Int J Mol Sci 23 (2022) 6769. 10.3390/ijms23126769.

[29] J.H. Garcia, E.A. Akins, S. Jain, K.J. Wolf, J. Zhang, N. Choudhary, M. Lad, P. Shukla, J. Rios, K. Seo, S.A. Gill, W.H. Carson, L.R. Carrete, A.C. Zheng, D.R. Raleigh, S. Kumar, M.K. Aghi, Multi-omic screening of invasive GBM cells reveals targetable transsulfuration pathway alterations, J Clin Invest (2023) e170397. 10.1172/JCI170397.

[30] Y. Hu, L. Wang, X. Han, Y. Zhou, T. Zhang, L. Wang, T. Hong, W. Zhang, X.-X. Guo, J. Sun, Y. Qi, J. Yu, H. Liu, F. Wu, Discovery of a Bioactive Inhibitor with a New Scaffold for Cystathionine γ-Lyase, J Med Chem 62 (2019) 1677–1683. 10.1021/acs.jmedchem.8b01720.

[31] K. Liu, Y. Abouelhassan, Y. Zhang, S. Jin, R.W. Huigens Iii, Transcript Profiling of Nitroxoline-Treated Biofilms Shows Rapid Up-regulation of Iron Acquisition Gene Clusters, ACS Infect Dis 8 (2022) 1594–1605. 10.1021/acsinfecdis.2c00206.

[32] N.A. Wolff, A.J. Ghio, L.M. Garrick, M.D. Garrick, L. Zhao, R.A. Fenton, F. Thévenod, Evidence for mitochondrial localization of divalent metal transporter 1 (DMT1), FASEB J 28 (2014) 2134–2145. 10.1096/fj.13-240564.

[33] D. Leitsch, A.L. Mbouaka, M. Köhsler, N. Müller, J. Walochnik, An unusual thioredoxin system in the facultative parasite Acanthamoeba castellanii, Cell Mol Life Sci 78 (2021) 3673–3689. 10.1007/s00018-021-03786-x.

[34] Y. Aoyagi, Y. Hayashi, Y. Harada, K. Choi, N. Matsunuma, D. Sadato, Y. Maemoto, A. Ito, S. Yanagi, D.T. Starczynowski, H. Harada, Mitochondrial Fragmentation Triggers Ineffective Hematopoiesis in Myelodysplastic Syndromes, Cancer Discov 12 (2022) 250–269. 10.1158/2159-8290.CD-21-0032.

[35] F. Deschner, T. Risch, C. Baier, D. Schlüter, J. Herrmann, R. Müller, Nitroxoline resistance is associated with significant fitness loss and diminishes in vivo virulence of Escherichia coli, Microbiol Spectr 12 (2024) e0307923. 10.1128/spectrum.03079-23.

[36] A.A. Johnson, T.L. Cuellar, Glycine and aging: Evidence and mechanisms, Ageing Res Rev 87 (2023) 101922. 10.1016/j.arr.2023.101922.

[37] A.A. Parkhitko, P. Jouandin, S.E. Mohr, N. Perrimon, Methionine metabolism and methyltransferases in the regulation of aging and lifespan extension across species, Aging Cell 18 (2019) e13034. 10.1111/acel.13034.

[38] Y. Ouyang, Q. Wu, J. Li, S. Sun, S. Sun, S-adenosylmethionine: A metabolite critical to the regulation of autophagy, Cell Prolif 53 (2020) e12891. 10.1111/cpr.12891.

[39] S. Laxman, B.M. Sutter, X. Wu, S. Kumar, X. Guo, D.C. Trudgian, H. Mirzaei, B.P. Tu, Sulfur amino acids regulate translational capacity and metabolic homeostasis through modulation of tRNA thiolation, Cell 154 (2013) 416–429. 10.1016/j.cell.2013.06.043.

[40] O.R. Salazar, P. N Arun, G. Cui, L.K. Bay, M.J.H. van Oppen, N.S. Webster, M. Aranda, The coral Acropora loripes genome reveals an alternative pathway for cysteine biosynthesis in animals, Sci Adv 8 (2022) eabq0304. 10.1126/sciadv.abq0304.

[41] R.H. Lampe, T.H. Coale, K.O. Forsch, L.J. Jabre, S. Kekuewa, E.M. Bertrand, A. Horák, M. Oborník, A.J. Rabines, E. Rowland, H. Zheng, A.J. Andersson, K.A. Barbeau, A.E. Allen, Short-term acidification promotes diverse iron acquisition and conservation mechanisms in upwelling-associated phytoplankton, Nat Commun 14 (2023) 7215. 10.1038/s41467-023-42949-1.

[42] N. Chondrogianni, I. Petropoulos, S. Grimm, K. Georgila, B. Catalgol, B. Friguet, T. Grune, E.S. Gonos, Protein damage, repair and proteolysis, Mol Aspects Med 35 (2014) 1–71. 10.1016/j.mam.2012.09.001.

[43] L.S. Tain, C. Jain, T. Nespital, J. Froehlich, Y. Hinze, S. Grönke, L. Partridge, Longevity in response to lowered insulin signaling requires glycine N-methyltransferase-dependent spermidine production, Aging Cell 19 (2020) e13043. 10.1111/acel.13043.

[44] M. Wang, X. Liang, M. Cheng, L. Yang, H. Liu, X. Wang, N. Sai, X. Zhang, Homocysteine enhances neural stem cell autophagy in in vivo and in vitro model of ischemic stroke, Cell Death Dis 10 (2019) 561. 10.1038/s41419-019-1798-4.

[45] E. Desideri, G. Filomeni, M.R. Ciriolo, Glutathione participates in the modulation of starvation-induced autophagy in carcinoma cells, Autophagy 8 (2012) 1769–1781. 10.4161/auto.22037.

[46] B.D. Paul, J.I. Sbodio, S.H. Snyder, Cysteine Metabolism in Neuronal Redox Homeostasis, Trends Pharmacol Sci 39 (2018) 513–524. 10.1016/j.tips.2018.02.007.

[47] X. Zhang, Q. Zhao, T. Wang, Q. Long, Y. Sun, L. Jiao, M. Gullerova, DNA damage response, a double-edged sword for vascular aging, Ageing Res Rev 92 (2023) 102137. 10.1016/j.arr.2023.102137.

[48] D. Menolfi, S. Zha, ATM, ATR and DNA-PKcs kinases-the lessons from the mouse models: inhibition ≠ deletion, Cell Biosci 10 (2020) 8. 10.1186/s13578-020-0376-x.

[49] C. McCarthy-Leo, F. Darwiche, M.A. Tainsky, DNA Repair Mechanisms, Protein Interactions and Therapeutic Targeting of the MRN Complex, Cancers (Basel) 14 (2022) 5278. 10.3390/cancers14215278.

[50] A. Ciccia, S.J. Elledge, The DNA damage response: making it safe to play with knives, Mol Cell 40 (2010) 179–204. 10.1016/j.molcel.2010.09.019.

[51] L. Paulet, A. Trecourt, A. Leary, J. Peron, F. Descotes, M. Devouassoux-Shisheboran, K. Leroy, B. You, J. Lopez, Cracking the homologous recombination deficiency code: how to identify responders to PARP inhibitors, Eur J Cancer 166 (2022) 87–99. 10.1016/j.ejca.2022.01.037.

[52] C. Pelletier, P. Prognon, P. Bourlioux, Roles of divalent cations and pH in mechanism of action of nitroxoline against Escherichia coli strains, Antimicrob Agents Chemother 39 (1995) 707– 713. 10.1128/AAC.39.3.707.

[53] P. He, S. Huang, R. Wang, Y. Yang, S. Yang, Y. Wang, M. Qi, J. Li, X. Liu, X. Zhang, M. Feng, Novel nitroxoline derivative combating resistant bacterial infections through outer membrane disruption and competitive NDM-1 inhibition, Emerg Microbes Infect 13 (2024) 2294854. 10.1080/22221751.2023.2294854.

